# Phagosome resolution regenerates lysosomes and maintains the degradative capacity in phagocytes

**DOI:** 10.1101/2020.05.14.094722

**Authors:** Charlene Lancaster, Aaron Fountain, Roaya M. Dayam, Elliott Somerville, Javal Sheth, Vanessa Jacobelli, Alex Somerville, Mauricio Terebiznik, Roberto J. Botelho

## Abstract

Phagocytes engulf unwanted particles into phagosomes that then fuse with lysosomes to degrade the enclosed particles. Ultimately, phagosomes must be recycled to help recover membrane resources that were consumed during phagocytosis and phagosome maturation, a process referred to as phagosome resolution. Little is known about phagosome resolution, which may proceed through exocytosis or membrane fission. Here, we show that bacteria-containing phagolysosomes in macrophages undergo fragmentation through vesicle budding, tubulation, and constriction. Phagosome fragmentation requires cargo degradation, the actin and microtubule cytoskeletons, and clathrin. We provide evidence that lysosome reformation occurs during phagosome resolution since the majority of phagosome-derived vesicles displayed lysosomal properties. Importantly, we show that clathrin-dependent phagosome resolution is important to maintain the degradative capacity of macrophages challenged with two waves of phagocytosis. Overall, our work suggests that phagosome resolution contributes to lysosome recovery and to maintain the degradative power of macrophages to handle multiple waves of phagocytosis.

**Summary:** Phagocytes engulf particles into phagolysosomes for degradation. However, the ultimate fate of phagolysosomes is undefined. Lancaster, Fountain et al. show that phagosomes fragment to reform lysosomes in a clathrin-dependent manner. This process helps maintain the degradative capacity of phagocytes for subsequent rounds of phagocytosis.

## Introduction

Phagocytosis is the receptor-mediated engulfment and sequestration of particulate matter into phagosomes and plays a critical role in infection clearance and tissue homeostasis (Gray and Botelho, 2017; Levin et al., 2016; Henson, 2017; Lancaster et al., 2019). After formation, the innocuous nascent phagosomes are converted into highly degradative phagolysosomes by fusing with lysosomes, leading to the cargo digestion (Gray and Botelho, 2017; Levin et al., 2016; Fairn and Grinstein, 2012; Pauwels et al., 2017); herein, we will use the term *lysosomes* to include a spectrum of late endosomes, terminal storage lysosomes, and endolysosomes, which are late endosome-terminal storage lysosome hybrids (Bright et al., 2016; Bissig et al., 2017). The phagolysosome thus acquires lysosomal membrane proteins like lysosomal-associated membrane proteins 1 and 2 (LAMP1 and LAMP2), a myriad collection of hydrolytic enzymes, and a luminal pH <5 established by the V-ATPase (Gray and Botelho, 2017; Levin et al., 2016; Kinchen and Ravichandran, 2010; Naufer et al., 2018; Kim et al., 2014; Garg et al., 2011; Levin-Konigsberg et al., 2019; Levin et al., 2017). This ensures digestion of particulates into its primordial components like amino acids and sugars, which are exported via transporters such as SLC-36.1 (at least in *Caenorhabditis elegans*) or in bulk via vesicular-tubular intermediates (Mantegazza et al., 2014; Gan et al., 2019).

The final stage of the life cycle of a phagosome is now referred to as phagosome resolution (Levin et al., 2016; Gray and Botelho, 2017). In unicellular eukaryotes like amoeba, phagosomes containing indigestible material are resolved via exocytosis, or egestion, expelling indigestible content and recycling the plasma membrane (Gotthardt et al., 2002; Stewart and Weisman, 1972). While this was assumed to occur in mammalian cells, recent work suggest that phagosomes undergo shrinkage and fission instead (Krajcovic et al., 2013; Krishna et al., 2016; Levin-Konigsberg et al., 2019). Partly, this occurs via processes regulated by the mechanistic Target of Rapamycin Complex-1 (mTORC1) and the lipid kinase PIKfyve (Krajcovic et al., 2013; Krishna et al., 2016). In addition, phosphatidylinositol-4-phosphate [PtdIns(4)P] on phagolysosomes recruits SKIP/PLEKHM2 to secure kinesin motors to lysosomes via the Arl8b GTPase, which collectively drive membrane extrusion via phagosome tubules, aiding in phagosome resolution. Interestingly, ER-phagosome contact sites formed via Rab7 and the oxysterol-binding protein-related protein 1L (ORP1L) transfer phagosomal PtdIns(4)P to the ER for turnover and elimination of PtdIns(4)P from resolving phagolysosomes (Levin-Konigsberg et al., 2019).

Surprisingly, little else is known about phagosome resolution, including how the phagosomal membrane is recycled, or how phagocytes retune their endomembrane state after degradation of phagosomal contents. In particular, while phagocytosis activates the transcription factor EB (TFEB) to induce expression of lysosomal proteins, boosting lysosome and cytoprotective functions (Gray et al., 2016; Visvikis et al., 2014), phagosomes consume “free” lysosomes during maturation, though we are not aware of a study that tested this. For long-lived macrophages (van Furth and Cohn, 1968; Parihar et al., 2010), “free” lysosomes and other membranes must be replenished to allow these cells to degrade multiple rounds of internalized cargo over their life, though it is unknown if and how lysosomes are reformed after phagosome maturation.

In this study, we report that phagosomes in macrophages undergo fission, tubulation, and splitting, culminating in phagosome fragmentation to reform lysosome-like organelles. We show that phagosome fragmentation required particle digestion, the actin and microtubule cytoskeleton systems, and clathrin. Moreover, we show that a first wave of phagocytosis reduced the number of “free” lysosomes and abated the degradative activity in phagosomes formed during a second round of phagocytosis. Importantly, given enough time, “free” lysosomes were replenished, and degradative activity within subsequent phagosomes was recovered in a clathrin- and biosynthetic-dependent manner. Overall, our study reveals a role of phagolysosome resolution in the reformation of lysosomes for reuse by macrophages to sustain multiple rounds of phagocytosis.

## Results

### Phagolysosomes with undigestible particulates are not egested

Despite numerous studies on phagosomal dynamics, the terminal fate of the phagolysosome remains obscure. For years, phagocytosis was assumed to end with the exocytosis of the phagolysosome and egestion of debris, a notion that likely arose from studies on phagotrophic protozoans that expel indigestible material by exocytosis (Clarke et al., 2002; Maniak, 2003; Stewart and Weisman, 1972; Gotthardt et al., 2002). However, more recent evidence suggests that mammalian phagosomes containing degradable cargo, like apoptotic cells or red blood cells, undergo shrinkage and/or fission (Krajcovic et al., 2013; Levin-Konigsberg et al., 2019). To investigate if phagolysosomes containing indigestible material are secreted in mammals, we allowed RAW 264.7 macrophages to internalize IgG-opsonized latex beads and followed the non-degradable cargo over 24 h by live-cell imaging. Over this period, we did not observe release of phagosome-sequestered beads (Fig. 1A-B and Video 1). Additionally, ionomycin, a calcium ionophore that induces exocytosis (Jans et al., 2004; Xu et al., 2013), did not cause the egestion of beads from macrophages (Sup. Fig. S1A and S1B, and Video 2), despite activating phospholipase C (Botelho et al., 2000b; Sup. Fig. S1C). Altogether, phagosomes containing non-degradable beads are retained inside mammalian macrophages instead of undergoing exocytosis.

**Figure 1.**
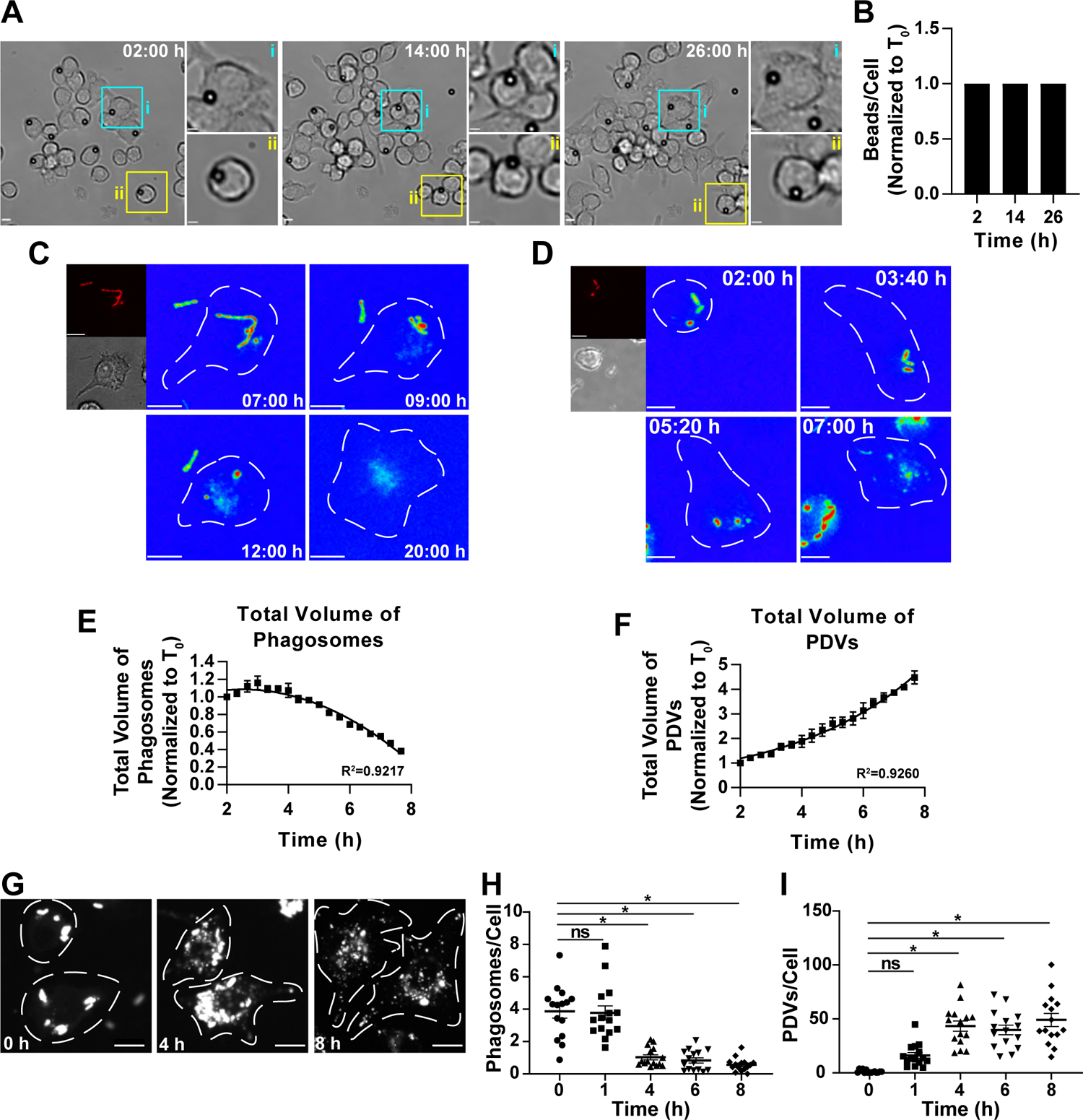
Phagolysosomes undergo fragmentation instead of exocytosis. **A.** RAW macrophages engulfed IgG-opsonized beads and were tracked for 24 h. Shown are DIC images. The coloured boxes indicate the areas enlarged and shown in *i* and *ii*. **B.** The number of latex beads within macrophages after indicated chase period and normalized to T_0_. Data shown as mean ± SEM from three independent experiments, 8 cells were quantified per independent experiment. **C, D** and **G.** RAW macrophages were challenged with IgG-opsonized, filamentous mCherry-*Lp* for 7 h (C) or with IgG-opsonized mRFP1-*E. coli* for 2 h (D, G) and then imaged live. White dashed lines indicate the boundary of the cell. C and D: The main panels show the red channel in a rainbow scale, red and blue are the highest and lowest intensity levels, respectively. The smaller panels show the red and DIC channels for the first frame. **E** and **F.** The total volume of phagosomes and PDVs per cell over time is shown as mean ± SEM of 3 independent experiments, with 10 to 25 cells analyzed per experiment. Volume was normalized to the first measurement, T_0_. **H** and **I.** Quantification of the number of PDVs and intact phagosomes. Data are means ± SEM of 3 independent experiments, 15 images were quantified per time point. A one-way ANOVA with Tukey’s post-hoc test compared each time point against the T_0_ (*, p < 0.05; ns, not significant). See videos 1, 3 and 4 for corresponding movies. Scale bars are 10 µm in main panels and 5 µm in insets.

### Phagolysosomes containing bacterial cargo undergo fragmentation

Recently, efferosomes and red blood cell-containing phagosomes were observed to undergo shrinkage and fragmentation (Krajcovic et al., 2013; Levin-Konigsberg et al., 2019). As these phagocytic targets are non-inflammatory, we thus queried whether phagosomes containing bacteria, which provoke an inflammatory response, would suffer a similar fate. We first utilized paraformaldehyde (PFA)-killed filamentous *Legionella pneumophila* (herein *Legionella* or Lp) expressing mCherry to track large phagosomes by live-cell imaging (Prashar et al., 2013). We used mCherry fluorescence as a proxy for the fate of phagosomes. Interestingly, within 7-20 h, the bacterial filaments collapsed into spheroidal bodies and there was a gradual increase in cytoplasmic puncta labelled with mCherry (Figure 1C and Video 3).

We further investigated this phenomenon by following PFA-fixed mRFP1-*Escherichia coli (E. coli)*-containing phagosomes (Figure 1D, Video 4) because *E. coli* are quasi-homogenous in shape and mRFP1 is resistant to degradation (Katayama et al., 2008). Using fluorescence intensity and volume thresholding to differentiate between phagosomes and phagosome-derived puncta, we calculated the total volume of phagosomes and the total volume of phagosome-derived vesicles (PDVs) per cell over time (details in Materials and Methods).

The total volume of phagosomes per macrophage decayed quadratically, while the total volume of PDVs increased exponentially over time, suggesting correspondence in the two phenomena (Figure 1E, 1F). These observations were complemented through fixed-cell, population-based assays, where we found that the number of intact *E. coli* remaining within macrophages decreased over time, while the number of puncta increased (Figure 1G, 1H, and 1I).

To assess if phagosomes fragmented even when they carry indigestible material, we prepared a semi-degradable cargo by aggregating fluorescently-labelled 100 nm polystyrene nanobeads using IgG. We presumed that these bead clumps could be broken apart through IgG proteolysis within the degradative phagolysosome. Indeed, internalized bead clumps gradually formed a cloud of fluorescence or smaller puncta, suggesting that these phagosomes split into smaller components (Supplemental Figure S1D, Video 5). Thereby, these experiments reveal that phagolysosomes containing indigestible, yet modular particulate cargo, can undergo fragmentation.

Finally, while wholesale phagosomes do not appear to exocytose, we assessed if phagosome-derived content could be secreted. We fed mRFP1-*E. coli* to macrophages for 1 h and chased for an additional 1, 6, or 24 h before collecting media and cell lysates. Probing cell lysates by Western blotting, we observed mostly intact mRFP1 at 1 h post-phagocytosis and the accumulation of a cleaved mRFP1 product at 6 and 24 h post-phagocytosis, which likely originated as a by-product of hydrolytic enzymes in phagolysosomes (Sup. Fig. S1E, S1F). We then evaluated the presence of mRFP1 and GADPH in cell media, the latter used to control for macrophage cell death. At 1 h post-phagocytosis, there was little mRFP1 in cell media, but at 6 and 24 h, cleaved mRFP1 was observable. By normalizing to GADPH, cleaved-mRFP1 appears to preferentially accumulate in media at 6 h vs. 1 h relative to GADPH, suggesting that phagosomal cargo can be secreted (Sup. Fig. S1E, S1F).

### Phagolysosome fragmentation is dependent on cargo degradation and the cytoskeleton

We next sought to understand some of the prerequisites in phagosome resolution. First, we hypothesized that phagosome resolution requires cargo degradation. To test this, we used either a protease inhibitor cocktail or NH_4_Cl/ concanamycin A (Con A) mixture to alkalinize the lysosomal pH (Naufer et al., 2018; Li et al., 2013). Vehicle-treated cells increased their PDV volume 4 h after phagocytosis relative to 1 h post-phagocytosis, consistent with phagosome fragmentation (Figure 2A-D). In comparison, cells treated with the protease cocktail or the NH_4_Cl/ConA mixture displayed lower production of PDVs 4 h after phagocytosis relative to the corresponding control cells (Figure 2A-D). Thus, cargo degradation is necessary for the fragmentation of the phagolysosome.

**Figure 2:**
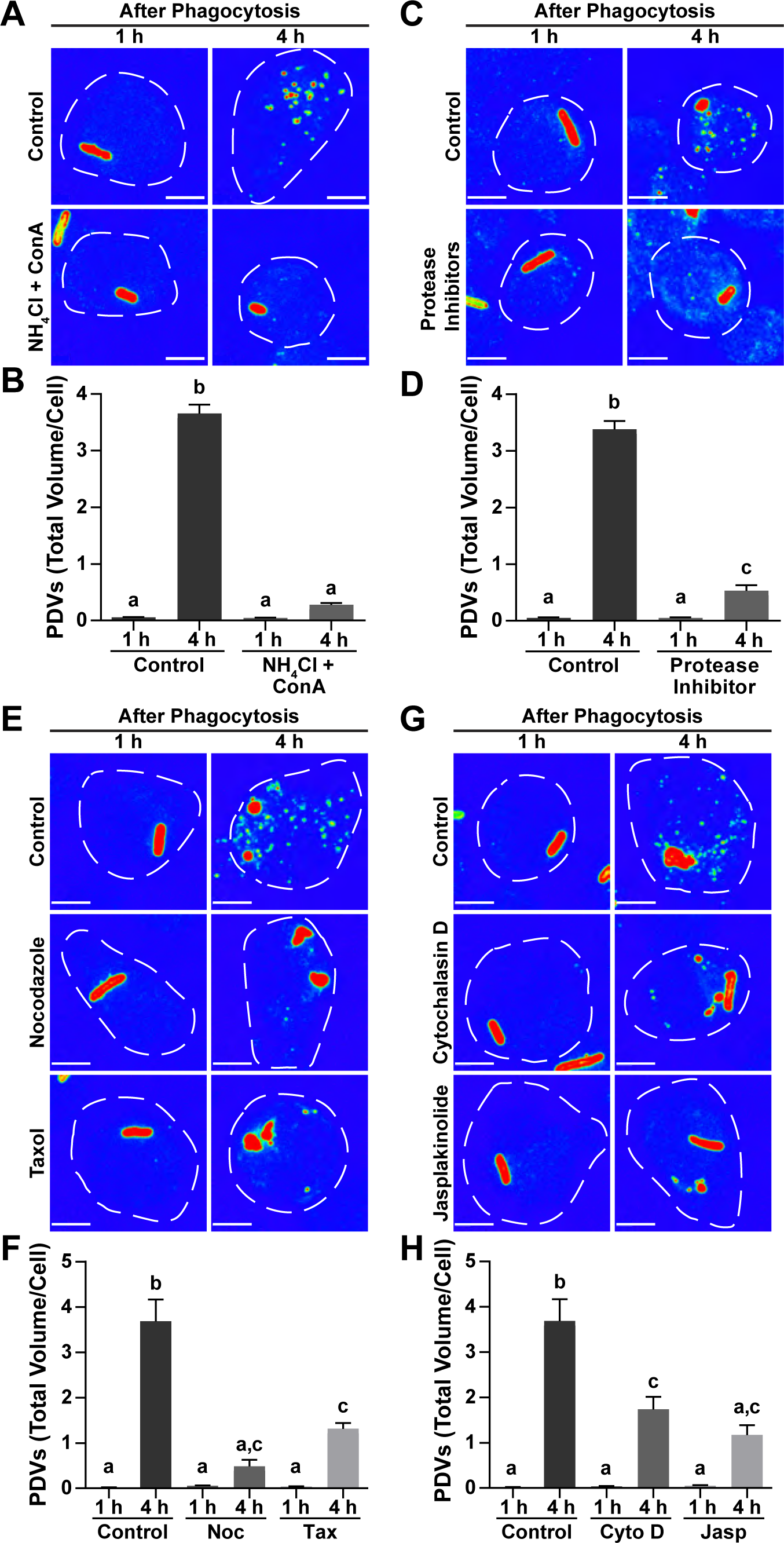
Fragmentation of the phagolysosome requires cargo degradation and the cytoskeleton. **A, C, E** and **G.** RAW macrophages engulfed *E. coli* for 15 min, and then chased for 30 min before treatment with either concanamycin A (ConA) and NH_4_Cl (A), protease inhibitor cocktail (C), microtubule inhibitors (E), actin inhibitors (G), or vehicle control (DMSO). Cells were fixed after 1 or 4 h post-phagocytosis and stained with an anti-*E. coli* antibody. Anti-*E. coli* antibody fluorescence is displayed in a rainbow scale, where red and blue are the highest and lowest fluorescence intensity levels, respectively. White dotted lines are the cells contours. Scale bars are 5 μm. **B, D, F,** and **H.** The total volume of PDVs per cell. Data are mean ± SEM of 3 independent experiments from 25 cells were quantified per experiment per condition and compared by one-way ANOVA with Tukey’s post-hoc test. Conditions labeled with different letters are statistically different (p < 0.05).

We then assessed the role of the cytoskeleton in phagosome fragmentation since the cytoskeleton is implicated in membrane scission events like coat protein assembly, motor-driven constriction, and membrane tubulation (Damiani, 2003; Gautreau et al., 2014; Ripoll et al., 2018; Bezanilla et al., 2015). To assess this, RAW macrophages internalized mRFP1-*E. coli* for 40 min and were then treated with the actin and microtubule stabilizing drugs, jasplakinolide and taxol, respectively, or with cytochalasin D or nocodazole to respectively depolymerize actin and microtubule. Unlike control conditions, all these drugs significantly lowered PDV volume at 4 h post-phagocytosis (Figure 2E-H). Therefore, the actin and microtubule cytoskeletons are required for efficient fragmentation of the phagolysosome.

### Clathrin is necessary for resolution of the phagolysosome

We next studied the dynamics of phagosome fragmentation and observed fission events that generated vesicles (Figure 3A, 3A’, Videos 6 and 7), tubules that either scissioned or collapsed back into the original organelle (Figure 3B, Video 8), or constriction events that generated large fragments rather than smaller vesicles (Figure 3C, Video 9). This suggests multiple mechanisms of phagosome fragmentation.

**Figure 3.**
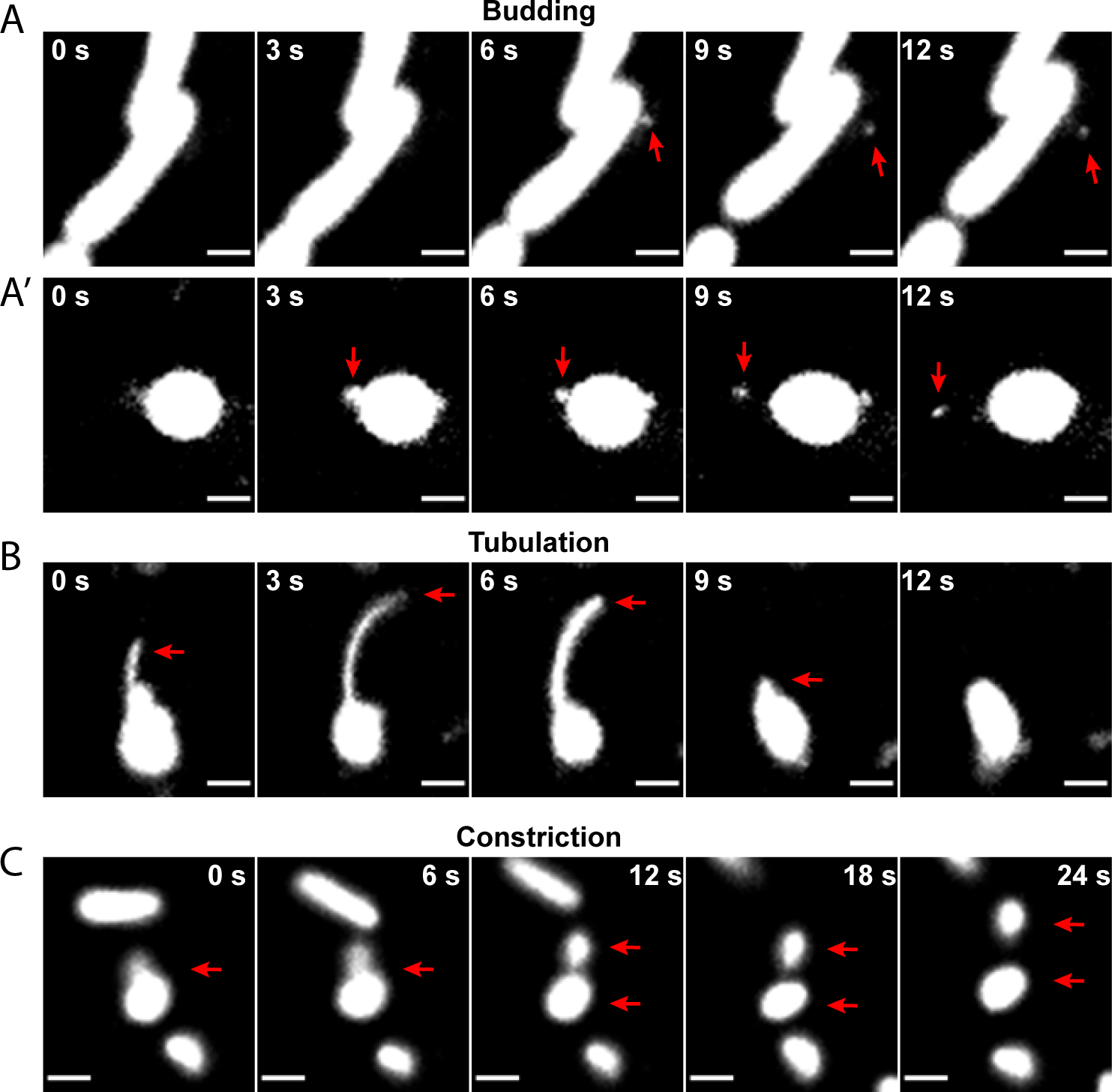
Phagosomes undergo different modes of fragmentation fission. RAW macrophages engulfed mRFP1*-E. coli* for 1 h, chased for 3 h to elicit phagosome fragmentation, and then imaged live for 10-min intervals at a frequency of 1 frame every 3 s. **A and A’.** Vesicle budding events in phagosomes were *E. coli* rods collapsed (A, early budding), or after initiating fragmentation (A’, late budding). **B** and **C.** Tubulation events producing small phagosomal fragments or retracting into the phagosome (B) and constrictions forming large phagosomal fragments (C). Red arrows track single events described above. See videos 6-9 for corresponding live-cell imaging. Scale bars are 2 µm.

Clathrin is an important mediator of vesicle budding and is well-known for its role at the plasma membrane during endocytosis and export from the *trans*-Golgi network (Kirchhausen et al., 2014). However, clathrin is also involved in budding events at the endosome, lysosome and autolysosomes (Stoorvogel et al., 1996; Traub et al., 1996; Rong et al., 2012). We thus investigated if clathrin is required for phagosome fragmentation. We first observed clathrin during phagosome maturation and resolution by live-cell imaging of RAW macrophages expressing a GFP-fusion of the clathrin-light chain (CLC-GFP). Our imaging analysis revealed clusters of clathrin localized in close proximity to phagolysosomes containing *Lp* (Figure 4A) or beads (Sup. Figure S2A). Quantifying the frequency of clathrin puncta associated with fission events was challenging, given the heterogeneity of phagosome resolution caused by varying phagosome age, the three-dimensionality of phagosomes, the ephemeral nature of clathrin puncta, and photobleaching. Yet, we observed that clathrin puncta co-occurred in 38 fission events out of 55, or about 70% of the fission events (Fig. 4B and 4C, Sup. Video 10). To assess if clathrin was bound to phagosomes, we isolated phagosomes and fixed them with PFA to lock protein complexes onto phagosomes before immuno-staining for LAMP1 and clathrin. We found clathrin-patches on phagolysosomes, indicating that clathrin is physically associated with these compartments (Figure 4D).

**Figure 4.**
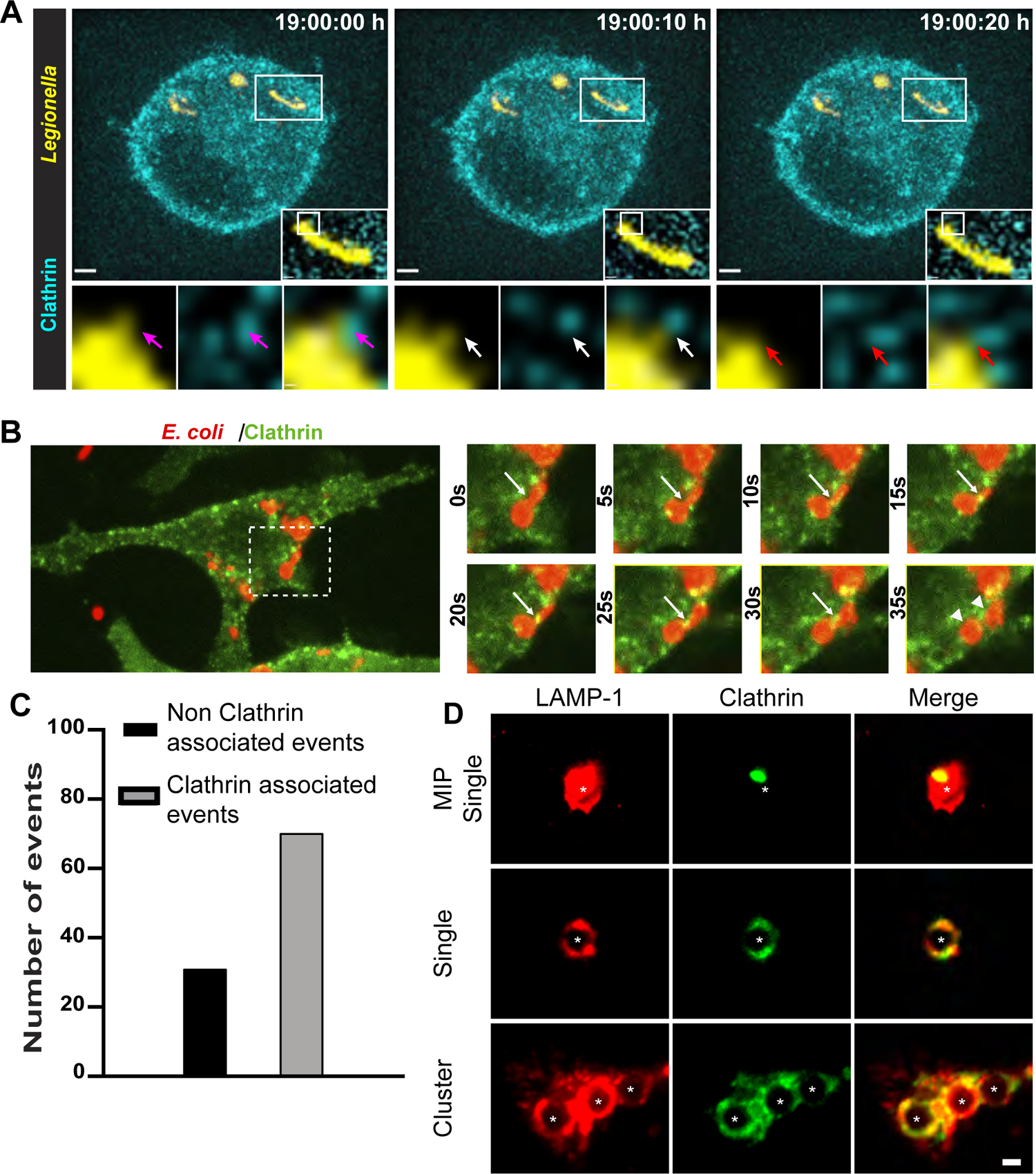
Clathrin localizes to the phagolysosome. **A** and **B.** Macrophages expressing clathrin light chain-GFP were allowed to internalized mCherry-*Lp* and chased overnight (A) or mRFP1-*E. coli* and chased for >1 h (B) before live cell imaging. For A, collapsed z-stacks are shown, while insets are deconvolved, single z-planes. White frames are enlarged areas in the inset and lower panels. Arrows show clathrin in close association with *Lp*. Scale bar: main panels are 2 μm; large insets are 0.5 μm; lower panels are 0.1 μm. For B, arrows show fission sites associated with a clathrin puncta, arrowheads show fragments post-fission. Images from Supplemental Video 10. **C.** Quantification of fission events associated with clathrin puncta**. D.** Isolated and fixed phagosomes (*) were immunolabeled for LAMP-1 (red) and clathrin (green). Top row: a z-stack of a single phagosome projected as the Maximum Intensity Projection, associated with a clathrin patch. Middle row: single-plane image of isolated phagosome showing clathrin patches. Bottom row: cluster of phagosomes co-isolated. Images are representative of 24 phagosomes analyzed. Scale bar is 2 µm.

We then evaluated the role of clathrin in phagosome fragmentation. To avoid interference with phagocytosis or phagosome maturation that might compromise phagosome resolution, we inhibited clathrin in macrophages with two inhibitors, Pitstop 2 and ikarugamycin, after 40-60 min of phagosome maturation (Von Kleist et al., 2011; Elkin et al., 2016). Using live-cell imaging of *Lp*-containing phagosomes over 13 h post-phagocytosis, we found that phagolysosomes in Pitstop-treated cells did not collapse to the same extent as in control cells, nor was there an increase in the volume of PDVs over many hours post-phagocytosis (Figure 5A, 5B and Video 11). Similarly, we observed a significant reduction in PDVs in Pitstop and ikarugamycin-treated cells, 4 to 8 h post-phagocytosis of *E. coli*, quantified by puncta volume (Figure 5C and 5D) and particle number (Sup. Figure S2B-E). Additionally, we used a rapamycin-induced dimerization system that acutely displaces clathrin to mitochondria (Robinson et al., 2010; Robinson and Hirst, 2013). Satisfyingly, this treatment impaired phagosome fragmentation relative to control conditions (Figure 5E, F). To determine that these observations were not constrained to RAW cells, we showed phagosome fragmentation in primary macrophages that engulfed mRFP1-*E. coli* and that this was impaired by Pitstop or ikarugamycin (Sup. Fig. S2F, G).

**Figure 5.**
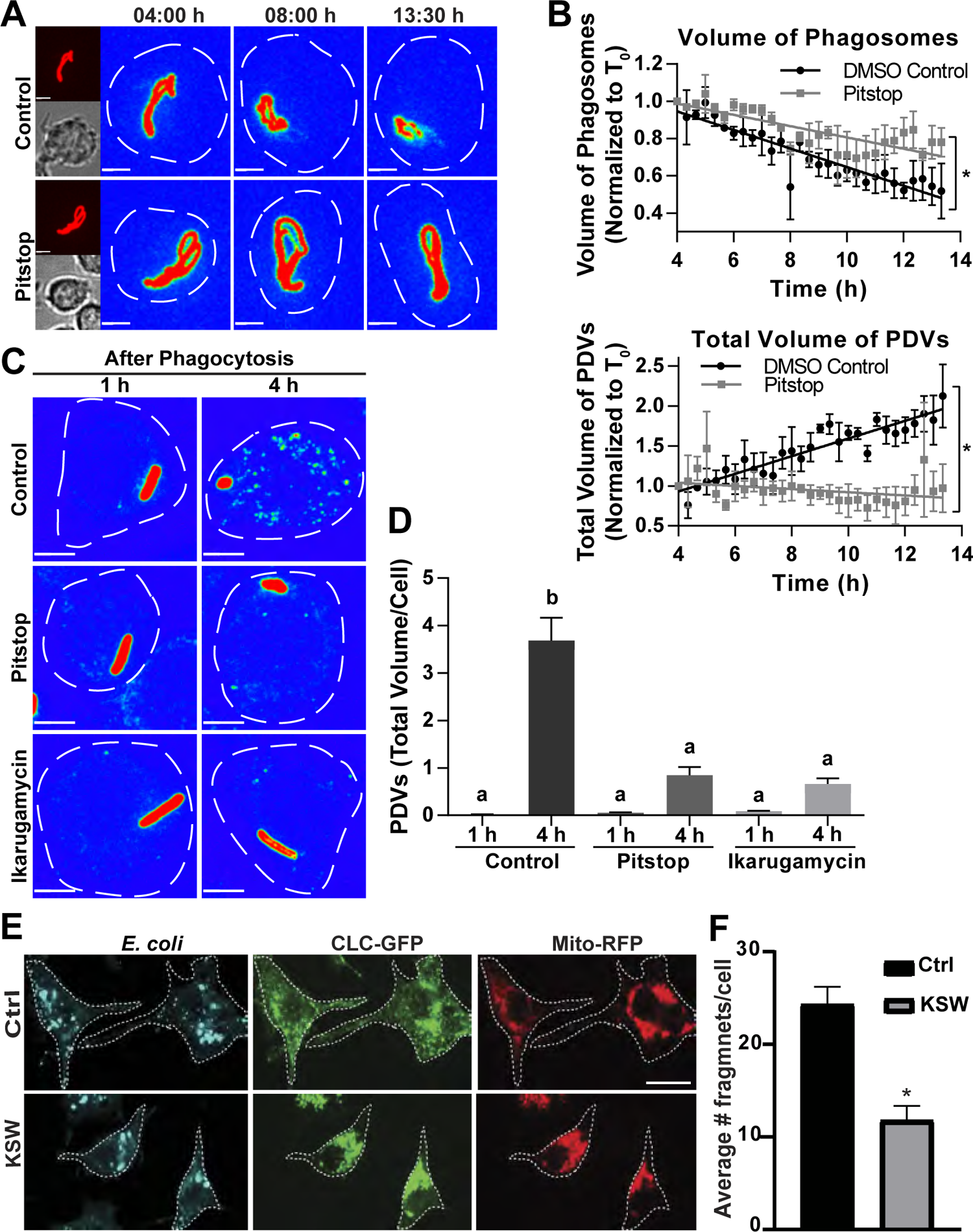
Clathrin is necessary for the resolution of the phagolysosome. **A.** Live cell imaging of RAW cells after phagocytosis of mCherry-*Lp*. After 4 h, Pitstop was added 10 min before imaging. **B.** Quantification of the volume of phagosomes and PDVs. Volumes were normalized to T_0_ for the respective treatment. Data are mean ± SEM of 3 independent experiments and * p < 0.05 indicates that the regressions are significantly different. **C**. RAW cells engulfed mCherry-*E. coli* for 15min, chased for 30 min, and then treated with either Pitstop or Ikarugamycin for 1 h or 4 h before fixation and imaging. Image representation is as described in Figure 2 legend. Scale bar: 5 µm. **D.** Volume of PDVs per cell for experiments shown in C. Data are means ± SEM of 3 independent experiments, 25 cells were quantified per experiment and results were tested by one-way ANOVA with Tukey’s post-hoc test. Different letters indicate results statistically different (p < 0.05). **E.** RAW macrophages expressing Mito-mCherry-FRB and GFP-FKBP-CLC were treated with rapamycin (KSW) or DMSO (Ctrl) and allowed to internalize *E. coli*. Cells were fixed 3 h post-phagocytosis, stained for *E. coli* and imaged. Scale bar: 10 µm. **F.** Number of fragments stained with anti-*E. coli* antibodies in control and rapamycin-treated cells. Data are represented as mean ± SEM of 3 independent experiments and statistically tested using an unpaired t-test (* p < 0.05).

The large GTPase dynamin is essential for clathrin budding, as it catalyzes the scission of clathrin-coated vesicles from cellular membranes (Mettlen et al., 2009). To assess dynamin involvement in the resolution of phagosomes, we utilized two dynamin inhibitors, dyngo-4a and dynole 34-2 (Hill et al., 2009; Mccluskey et al., 2013). Dynole 34-2 or dyngo-4a-treated cells produced a significantly smaller total volume of PDVs than control cells 4 h post-phagocytosis (Supplemental Figure S3). Collectively, the clathrin machinery mediates the resolution of the phagolysosome.

### Phagosome-derived vesicles have lysosomal characteristics

To investigate if PDVs retained a predominant lysosomal character, we looked at their association with endo-lysosomal markers. Macrophages were challenged with ZsGreen-*E. coli* for 6 h to elicit phagosome fragmentation, then fixed and immunostained against the lysosomal-associated membrane protein 1 (LAMP1) and LAMP2. Approximately 80% of PDVs (defined as ZsGreen-positive compartments with an area >0.1 µm^2^ but <4 µm^2^ to exclude the parental phagosomes) were positive for LAMP1 and LAMP2 (Figure 6A and 6B). Subsequently, we determined that > 80% of PDVs were acidic and proteolytically-active by respectively using LysoTracker Red, a membrane-permeable and acidotrophic fluorophore, and the Magic Red Cathepsin L fluorogenic substrate (Figure 6C-6F). Altogether, phagolysosome-derived compartments predominantly retain endo-lysosomal features of the mother phagolysosome.

**Figure 6.**
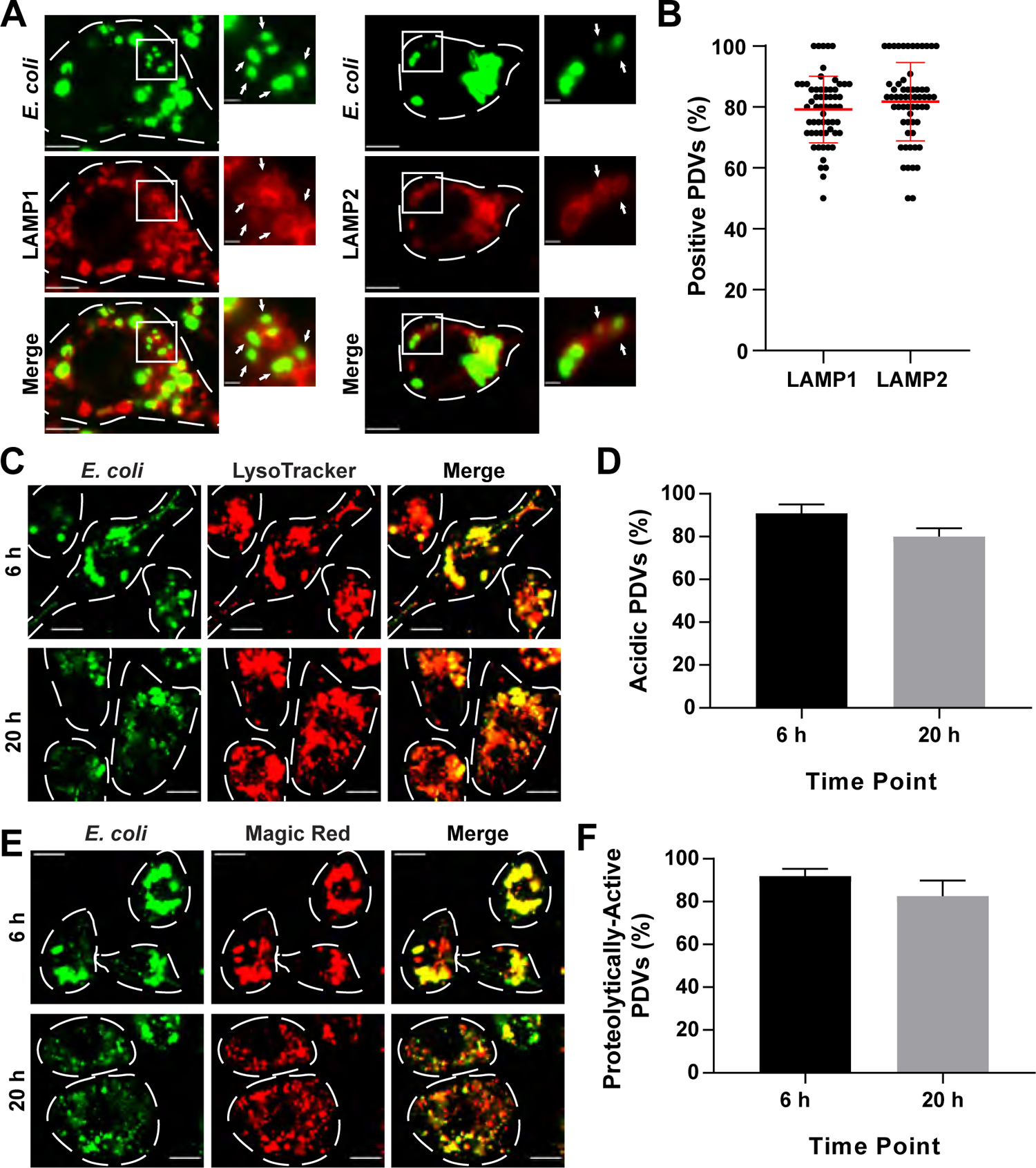
Phagosome-derived vesicles exhibit lysosomal properties. **A.** Macrophages were challenged with ZsGreen-*E. coli* for 6 h, fixed and immunostained for LAMP1 or LAMP2. White dotted lines are cell contours. Arrows indicate colocalization of LAMP1/2 to PDVs. Scale bars are main panels: 5 μm; insets: 1 μm for left and right panels, respectively. **B.** Percentage of PDVs positive for LAMP1/2. Data are mean ± SD of 60 cells from 3 independent experiments. **C** and **E.** PDVs from ZsGreen-*E. coli* phagosomes visualized at the 6 and 20 h post-phagocytosis, in cells stained with LysoTracker Red (C) or Magic Red cathepsin L substrate (E) for 1 h prior to live-cell imaging acquisition. Scale bar is 10 µm. **D** and **F.** PDVs were scored for co-occurrence with Lysotracker (C) or Magic Red (E). See methods for details. Data are mean ± SEM of 3 independent experiments. 25-110 cells were quantified at 6 or 20 h post-phagocytosis per experiment per time point analysed. Data was statistically analysed by unpaired t-test (* p < 0.05).

### Phagolysosome fragmentation reforms lysosomes

Phagosomes extensively fuse with lysosomes during maturation, consuming “free” lysosomes which may exhaust the degradative capacity of the phagocyte. Given macrophages long life-span and capacity to undertake successive rounds of phagocytosis (Cannon and Swanson, 1992; Parihar et al., 2010; van Furth and Cohn, 1968), we postulated that phagosome fragmentation may participate in lysosome reformation and maintaining the degradative capacity of macrophages.

To test this hypothesis, we assessed the number of LAMP1-positive puncta as a proxy for “free” lysosomes in resting macrophages or after phagocytosis of indigestible latex beads, precluding phagosome resolution. We saw significantly fewer “free” lysosomes in macrophages that engulfed latex beads and undertook phagosome maturation (60 min) relative to resting macrophages or those with phagosomes but that had limited time to fuse with lysosomes (15 min; Figure 7A and 7B). In comparison, there was no apparent difference in “free” lysosome number between resting macrophages and those with immature phagosomes (15 min), consistent with low phagosome-lysosome fusion and demonstrating that our quantification method was not affected by bead crowding in the cytoplasm (Figure 7A and 7B).

**Figure 7:**
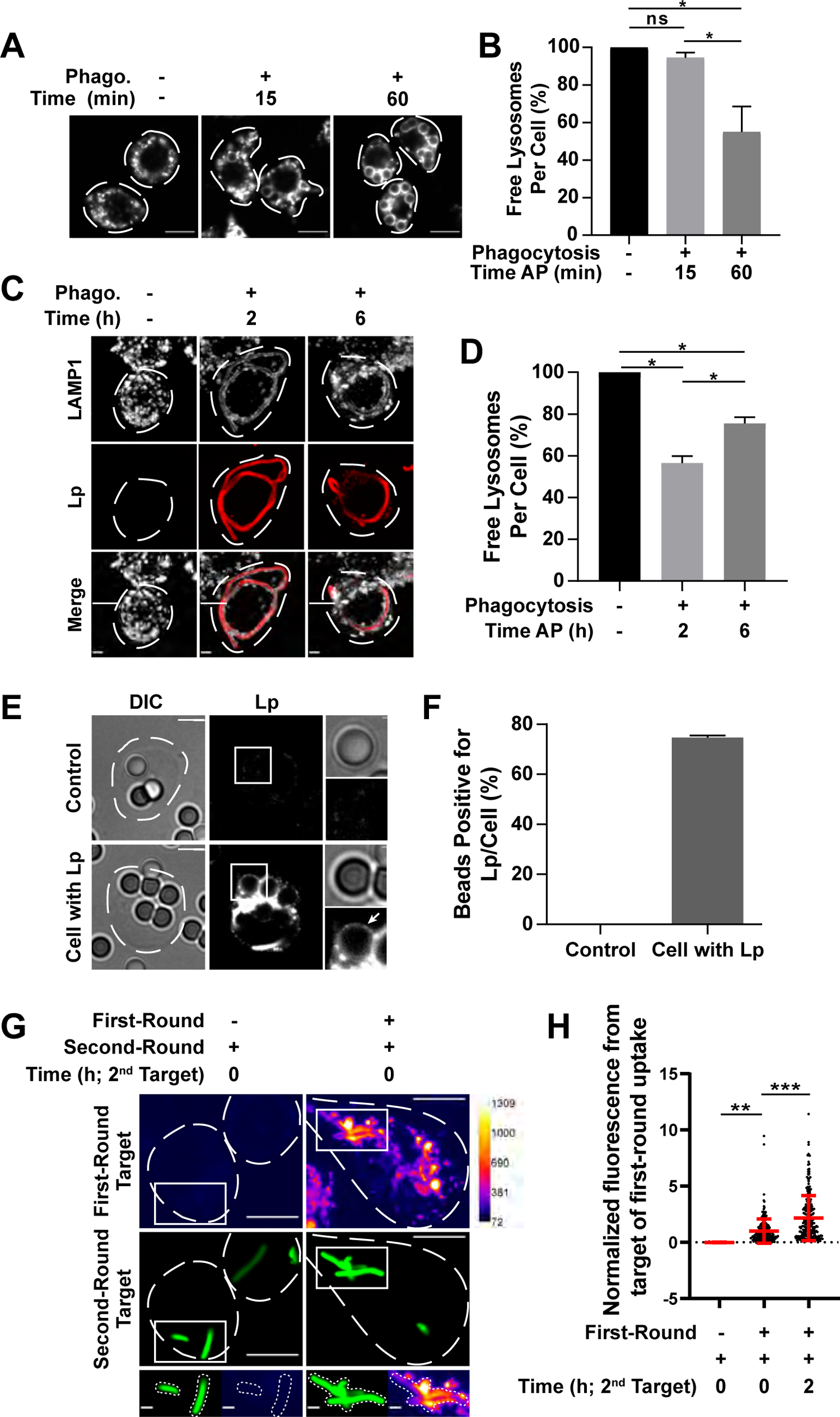
Phagosome maturation consumes free lysosomes. **A** and **C.** Macrophages were challenged with IgG-beads (A) or mCherry-*Lp* (C, Lp), and immunostained for LAMP1. Images show distribution of LAMP1 (A, C) and *Lp* (C). **B.** Quantification of the number of free lysosomes per cell for images described in A. **D.** Number of free lysosomes per cell in images described in C. Data were normalized to resting macrophages. For B and D: Data are means ± SEM of 3 independent experiments, where 60 cells (B) and 20 cells (D) were quantified per condition in each experiment, normalized to resting macrophages, and statistically tested as before. Scale bars are 10 µm and 5 µm in A and C, respectively. **E.** RAW cells that engulfed *Lp* were chased 6 h to allow for the formation of *Lp*-containing PDVs (bottom panel), and subsequently presented with IgG-opsonized beads for a second round of phagocytosis for 1 h. Cells were fixed and immunostained for *Lp*. The arrow shows bacterial debris in the bead phagosome. Scale bars are main panels: 5 µm; insets: 0.5 µm. **F.** The percentage of beads that was positive for *Lp* fragments in E. Data are mean ± SEM of 3 independent experiments. 20 cells were quantified for each condition per experiment. **G.** Macrophages were presented with mRFP1-*E. coli* for 7 h as a first round of phagocytosis before being challenged with ZsGreen-*E. coli* for a second round of uptake. As a control, resting macrophages were challenged only with one round of phagocytosis using ZsGreen-*E. coli*. Scale bars are main panels: 10 μm; insets: 2 μm. **H.** Quantification of mean mRFP1 fluorescence intensity (derived from first-round phagosomes) on ZsGreen-*E. coli*-containing phagosomes (second-round phagosomes) as described in G. Data was normalized to macrophages without the first-wave of phagocytosis and presented as mean ± SD of 148-348 cells across 3 independent experiments. Conditions were compared statistically using a one-way ANOVA with Tukey’s post-hoc test (**, p < 0.01; ***, p<0.001).

To determine if phagosome resolution aids in lysosome regeneration, we then employed *Lp*; these form extensive phagosomes that do not overcrowd the cytoplasm because of coiling, thus facilitating visualization of lysosome number. Free lysosomes were quantified by applying a mask on fluorescent *Lp* and quantifying the number of LAMP1-positive puncta outside of this mask. We compared resting cells and cells with *Lp*-containing phagosomes for 2 h or 6 h post-phagocytosis to respectively elicit phagosome-lysosome fusion (2 h) and fragmentation (6 h). As with the bead-containing phagosomes, cells that were incubated for 2 h post-phagocytosis suffered a 45% reduction in the number of “free” lysosomes (Figure 7C and 7D). Interestingly, and relative to 2 h post-phagocytosis, we observed a bounce in the number of “free” lysosomes after the onset of phagosomal fission (6 h; Figure 7C and 7D), suggesting that macrophages began to recover LAMP1-positive lysosomes through phagosomal fragmentation.

We next hypothesized that phagosome-derived lysosome-like compartments can fuse with subsequent phagosomes. To test this, we challenged macrophages with two rounds of phagocytosis. In one model, we allowed macrophages to engulf PFA-killed mCherry-expressing *Lp* followed by a 6 h chase to prompt phagosome resolution. We then challenged these macrophages to engulf latex beads followed by 1 h of phagosome maturation. We observed transfer of mCherry that originated from within *Lp* (first phagocytosis) onto bead-containing phagosomes (second phagocytosis; Figure 7E and 7F). In a second model, macrophages were first allowed to form phagosomes containing mRFP1-*E. coli* (first phagocytosis) for 1 h, and either immediately (0 h) or after 2 h to begin phagosome resolution, the cells were then challenged with ZsGreen-expressing *E. coli* (second phagocytosis) and chased for 1 h to elicit maturation. Using this model, macrophages that were given time to resolve the first phagosomes exhibited a higher degree of mRFP1 colocalization with secondary phagosomes than macrophages deprived of a chase time period (Figure 7G and 7H). Altogether, these observations suggest that PDVs fuse with secondary phagosomes, consistent with these being lysosome-like organelles.

### Clathrin-mediated phagosome fragmentation is required to recover degradative capacity

Since phagolysosomes consume lysosomes, we surmised that this may reduce the degradative capacity of phagosomes produced in subsequent rounds of phagocytosis. In turn, secondary phagosomes may exhibit lower degradative activity than their predecessors. To test this hypothesis, we challenged macrophages with two rounds of phagocytosis. In the first-round, macrophages engulfed either IgG-opsonized latex beads or *E. coli*, followed by a second phagocytic wave using IgG-opsonized beads coated with the fluorogenic protease substrate DQ-BSA. We observed that the fluorescence of DQ-BSA-coated beads was lower in those macrophages that internalized one prior round of beads or *E. coli* 1 h earlier, compared to macrophages with no prior phagocytosis (Figure 8A and 8B, red dots in mock primary vs. red dots in 1 h *E. coli* or 1 h beads). Importantly, the proteolytic activity of DQ-BSA phagosomes formed during the second-round of phagocytosis recovered significantly in macrophages chased for 6 h after initially engulfing *E. coli* (Figure 8A and 8B; blue vs. red dots in *E. coli* condition). Conversely, the fluorescence intensity associated with DQ-BSA beads was similar in cells that engulfed indigestible beads 1 h or 6 h prior during the first-round of uptake, though there was a trend upward at 6 h (Figure 8A and 8B, blue vs. red dots in primary bead condition).

**Figure 8.**
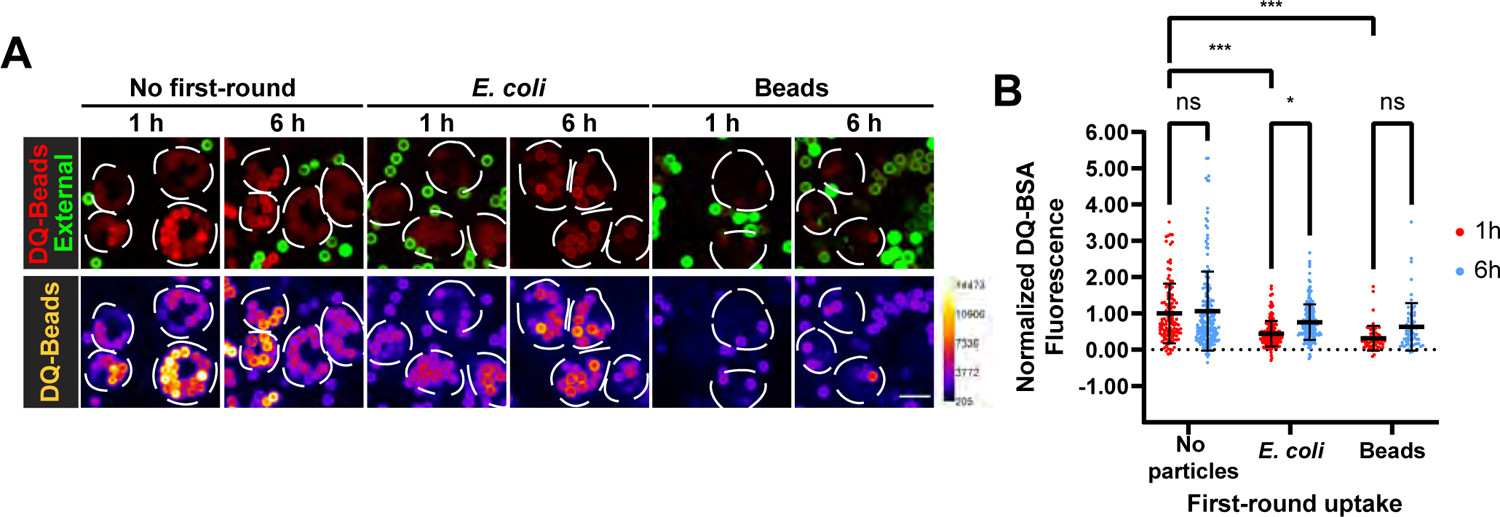
Phagosome resolution recovers degradative capacity in macrophages. **A** Macrophages either were given no particles or were allowed to engulf for 1 h unlabelled *E. coli* or unlabelled IgG-opsonized beads for the first round of phagocytosis, followed by 1h or 6 h to elicit maturation/resolution, before the addition of DQ-BSA-opsonized beads for a second-round of engulfment. External beads were immunostained (green) and cells imaged live. The lower panels show the red channel (DQ-BSA) in fire scale. Scale bars are 10 µm. **B.** Quantification of DQ-BSA fluorescence intensity of internalized DQ-BSA beads from experiments described in A. Intensity of each internal DQ-BSA bead was corrected by subtracting the mean intensity of external DQ-BSA beads. Data is presented as mean ± SD normalized to the mean of 1 h/vehicle condition and is based on 6 independent experiments, where 150-503 cells were quantified per condition (*, p < 0.05; **, p< 0.01; ***, p<0.001; ns, not significant).

Collectively, these observations indicate that the degradative capacity of macrophages decreases with phagocytosis but it is recovered most efficiently upon phagosome resolution.

We next tested whether clathrin-mediated phagosome resolution was required for macrophages to regain their degradative capacity. This required exposing macrophages to ikarugamycin for 6 h post-phagocytosis to elicit phagosome resolution. This prolonged treatment raised concerns of potential interference with basal macrophage functions that we assessed by determining the degradative activity of phagosomes on DQ-BSA-beads in macrophages exposed to clathrin inhibitors for 6 h. Indeed, the proteolytic activity associated with phagosomes in cells treated with ikarugamycin was reduced, relative to those exposed to vehicle for 1 h or 6 h (Figure 9A and 9B, no prior phagocytosis). This perturbation in phagosome-associated proteolytic activity likely represents an impairment in biosynthetic trafficking of proteases. We then examined the combined effect of clathrin inhibition and prior phagocytosis.

**Figure 9:**
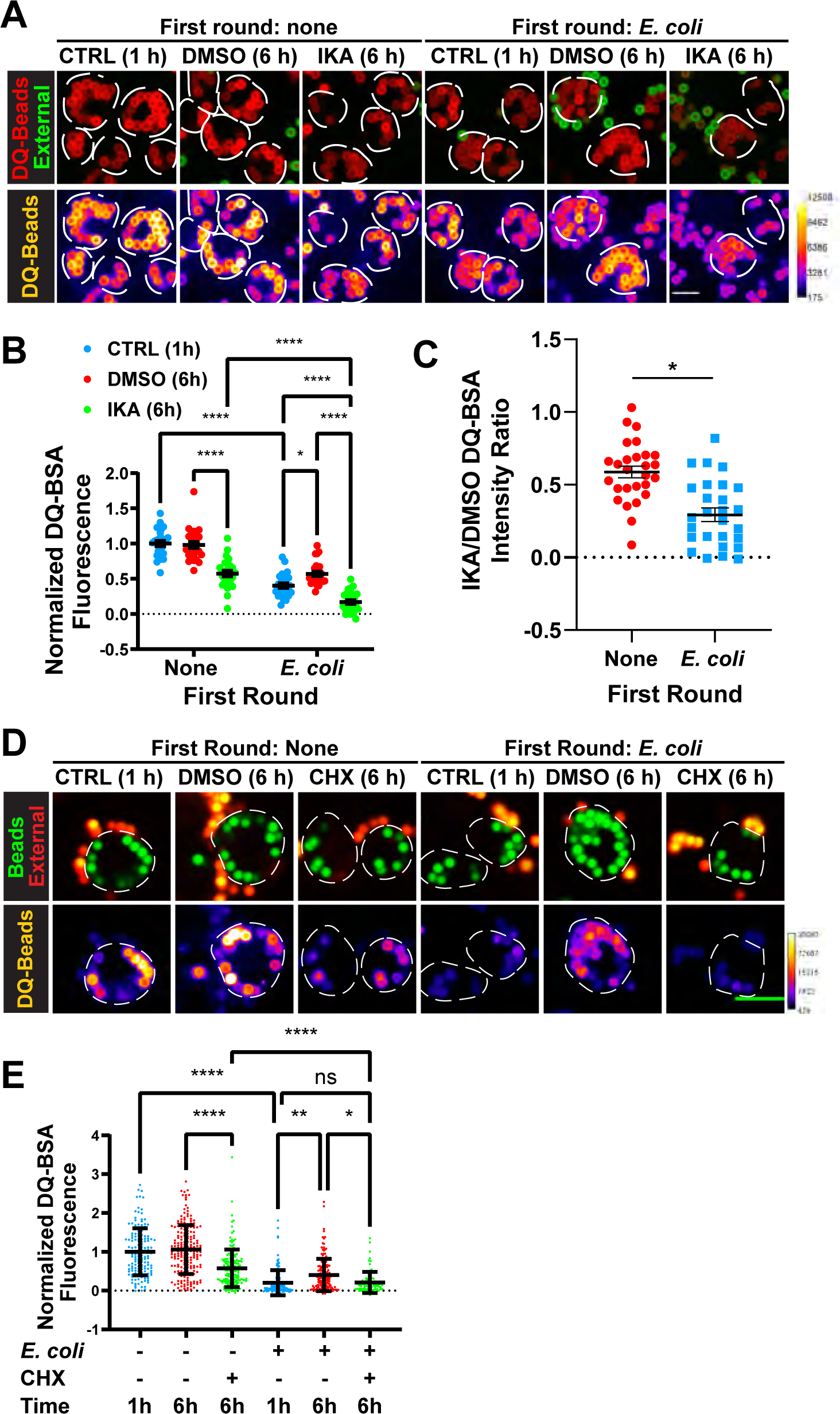
Clathrin-mediated phagosome resolution is required for degradative capacity of macrophages. **A** and **D.** Macrophages either did not receive a primary target or internalized *E. coli* for 1h, followed by 1h or 6 h chase before the addition of DQ-BSA-opsonized beads (red in A, green in D). External beads were immunostained (green in A, red in D, but appears yellow) and imaged live. The lower panels show the DQ-BSA fluorescence in fire scale. Scale bar: 10 µm. For A, macrophages were treated with the clathrin inhibitor, ikarugamycin (IKA), or vehicle (DMSO) 1 h after phagocytosis. For D, macrophages were treated with cycloheximide (CHX), or vehicle (DMSO) 1 h after phagocytosis. **B** and **E.** Quantification of mean DQ-BSA fluorescence intensity of internalized DQ-BSA beads from experiments described in A (B) and D (E) and processed as in Figure 8. For B, data is from 27 images across 3 independent experiments, with each image containing 1 to 8 cells; For E, data is based on 3 independent experiments, where 82-193 cells were quantified per condition. Conditions were compared statistically using a one-way ANOVA with Tukey’s post-hoc test for B, and two-way ANOVA with Tukey’s post-hoc test for E. **C.** Ratio of the mean fluorescence intensity of DQ-BSA in ikarugamycin-treated cells to DMSO-treated cells from data in B and shown as mean ± SEM. Statistical analysis was done using an unpaired t-test (*, p < 0.05; **, p< 0.01; ***, p<0.001; ns, not significant).

First, macrophages that engulfed *E. coli* 1 h prior had significantly lower DQ-BSA fluorescence in subsequent phagosomes than those macrophages without a prior round of uptake (Figure 9A and 9B, blue dots for cells with no *E. coli* vs. *E. coli* phagocytosis), consistent with consumption of lysosomes by the first wave of phagosomes. Second, there was a significant increase in DQ-BSA fluorescence in macrophages that engulfed *E. coli* 6 h prior, relative to those had engulfed *E. coli* 1 h prior (Figure 9A and 9B, blue and red dots under *E. coli* primary phagocytosis). This is consistent with results in Figure 8. Importantly, this recovery was abolished when cells were pre-treated with ikarugamycin (Figure 9B, green dots vs. blue and red dots under *E. coli* primary phagocytosis). We then ascertained if the combined effect of clathrin inhibition and phagocytosis was greater than clathrin inhibition alone, by normalizing the fluorescence of DQ-BSA-bead from ikarugamycin-treated cells to vehicle-treated cells. Our data show a significant abatement in this ratio for macrophages that undertook two rounds of phagocytosis relative to those that engulfed only DQ-BSA-beads (Figure 9C). Finally, we postulated that the biosynthetic pathway may aid in maintaining the degradative capacity macrophages during phagosome resolution.

First, we observed that cycloheximide did not impair phagosome fragmentation, suggesting the first wave of phagosomes had sufficient hydrolytic enzymes to complete digestion and resolution (Sup. Fig. S4A, B). However, there was a reduction in phagosome-associated proteolytic activity in cells treated with cycloheximide with or without a prior round of phagocytosis (Fig 9D, E).

Finally, we then examined the role of TFEB in recovering degradative capacity of macrophages by using RAW cells deleted for TFEB (Pastore et al., 2016). First, we observed no difference in the baseline degradation of DQ-BSA-bead phagosomes between wild-type and Tfeb^-/-^ cells with no prior phagocytosis (Sup. Fig. S4C, D). Moreover, after 1 h of phagocytosis with *E. coli*, there was a decreased in DQ-BSA-bead fluorescence in both wild-type and Tfeb^-/-^ macrophages (Supg. Fig. S4C, D). Importantly, the wild-type macrophages recovered some of this fluorescence level after 6 h post-phagocytosis of *E. coli* (Sup. Fig. S4C, D, compare red dots for *E. coli* 6 h vs. reds dots for no prior phagocytosis and *E. coli* 1 h). This recovery was partially abated in Tfeb^-/-^ macrophages relative to knockout cells with no prior phagocytosis or after 1 h of E. coli phagocytosis (Sup. Fig. S4C, D). However, we note that under the observed conditions, we did not observe a statistical difference between wild-type and Tfeb^-/-^ macrophages after 6 h of E. coli uptake (Sup. Fig. S4C, D). Overall, phagosome resolution is important to recycle lysosomes and maintain degradative proficiency of macrophages to handle subsequent rounds of phagocytosis. Phagosome resolution is coupled to biosynthesis and likely to lysosome adaptation pathways.

## Discussion

For every phagosome formed, membrane resources, including plasma membrane, endosomes, and lysosomes, are consumed and thus membrane resynthesis and retrieval is critical to keep phagocytes functional (Zent and Elliott, 2017). However, it is largely uncharacterized how phagocytes balance cellular resources, though synthesis of plasma membrane lipids has been observed (Werb and Cohn, 1972). Here, we show that macrophages predominantly resolve phagosomes via fragmentation, rather than through egestion. This fission process leans on the degradation, the cytoskeleton, clathrin and dynamin. Moreover, we present evidence that phagolysosome resolution is critical in restoring lysosomes that have been consumed during phagosome maturation and to sustain proteolytic activity in macrophages that ceaselessly iterate phagocytosis, with the biosynthetic pathway likely supplying new hydrolytic enzymes.

### The ultimate fate of phagolysosomes

It has been assumed that phagolysosomes undergo exocytosis to expel indigestible material and return membrane to the surface of the phagocyte (Stewart and Weisman, 1972; Maniak, 2003; Clarke et al., 2002; Gotthardt et al., 2002). However, we failed to observe exocytosis of phagosomes containing either bacteria or beads. Instead, we observed the occurrence of smaller vesicles derived from phagosomes, as well as the disappearance of recognizable phagosomes. This phagosome fragmentation proceeded through splitting, tubulation, and budding, suggesting at least three distinct mechanisms. This aligns with previous observations showing that efferosomes and phagosomes containing red blood cells shrink or fragment into phagosome-derived membrane compartments (Krajcovic et al., 2013; Levin-Konigsberg et al., 2019; Poirier et al., 2020). Collectively, these observations suggest that phagosome resolution is typically driven by fragmentation using distinct mechanisms of membrane fission rather than exocytosis, irrespective of the nature of the cargo. Yet, we have not excluded exocytosis as a mechanism of phagosome resolution for all modes of phagocytosis, such as the uptake of indigestible particulates captured by alveolar macrophages or resolution of phagosomes with pathogenic microorganisms (Ma et al., 2006; Alvarez and Casadevall, 2006). Moreover, some phagosome-derived cargo may eventually undergo secretion as detected by cleaved bacteria-derived mRFP1 in the media of phagocytes post-phagocytosis – this may occur through direct PDV secretion, or indirectly, through exchange with pre-existing compartments. This will need to be further explored.

In addition, while tubulation, budding, and splitting are detectable during phagosome resolution, it is important to state that membrane fission occurs throughout the lifetime of the phagosome, not simply during resolution (Saffi and Botelho, 2019). For example, COPI-mediated fission occurs on phagosomes to extract transferrin receptor from early phagosomes (Botelho et al., 2000a), while cytoskeleton mediates recycling from phagosomes enclosing latex beads (Damiani, 2003). Moreover, MHC-II::peptide complexes exit phagosomes likely before resolution ensues, though the exact mechanism and timing is unclear (Harding and Geuze, 1992; Mantegazza et al., 2013, 2014; Ramachandra et al., 1999). Finally, tubules can dynamically form between phagosomes to exchange phagosomal content with implications towards antigen presentation (Mantegazza et al., 2014). All these processes occur before phagosomes become unidentifiable as such. Thus, we propose a model in which membrane fission happens throughout the lifetime of the phagosome, where early and mid-maturation serves to sort and remove cargo from the phagosome without its disintegration, whereas end-of-life fission culminates in phagosome disappearance. It will be exciting to define if these represent distinct fission complexes or common fission assemblies that are regulated differently during the lifespan of the phagosome.

### Cargo degradation in phagolysosome resolution

Through inhibition of lysosomal proteases and neutralization of the phagosomal pH, we provide evidence that cargo degradation is required for phagosomal resolution. Mechanistically, we envision two non-mutually exclusive possibilities for this requirement. First, loss of physical integrity by cargo degradation may be a *sine qua non* for phagosome fission. Supporting this concept, undegradable large latex beads remain enclosed within phagosomes for prolonged periods of time. On the other hand, IgG-agglutinated nanobead clusters can fragment through degradation and disaggregation of IgG, which allows phagosomal fragmentation. Second, digestible cargo like bacteria, contains a complex mix of macromolecules that are degraded to release monomers like amino acids that become available for use within the phagocyte. This bolus of amino acids may be sensed by intraluminal amino acid sensors like the V-ATPase, or by cytoplasmic sensors like CASTOR, after amino acid transport across the phagolysosome membrane, locally promoting mTORC1 activation (Saxton et al., 2016; Chantranupong et al., 2016; Zoncu et al., 2011; Inpanathan and Botelho, 2019). While mTORC1 drives many anabolic processes like protein and lipid synthesis, mTORC1 also plays a role in coordinating phagosome and lysosome dynamics, including efferosome fission (Krajcovic et al., 2013; Inpanathan and Botelho, 2019; Saric et al., 2016; Hipolito et al., 2019). Thus, cargo degradation coupled to mTORC1 activation could help orchestrate phagosome resolution. This would parallel the role of mTORC1 re-activation during degradation of autophagic cargo to regenerate lysosomes (Yu et al., 2010).

### Requirements for phagosome resolution

We observed that drugs that stabilized or depolymerized the actin and microtubule cytoskeletons, and inhibitors of clathrin and dynamin, all impaired phagosome fragmentation. These, together with our observations that phagosomes suffer splitting, tubulation, and budding, implies that phagosome resolution depends on multiple mechanisms. Actin may play a role in membrane constriction via acto-myosin complexes that could distort phagosomes (Liebl and Griffiths, 2009; Curchoe and Manor, 2017; Poirier et al., 2020). Alternatively, short branched actin assemblies are associated with fission of endosome/lysosomes through contacts with the endoplasmic reticulum (Rowland et al., 2014; Hoyer et al., 2018). For example, the actin-nucleating protein Wiskott–Aldrich syndrome protein and SCAR homologue (WASH) and FAM21 are recruited to sites of membrane tubulation and fission at ER-endosome contact sites (Dong et al., 2016; Rowland et al., 2014; Hoyer et al., 2018). This could also be coupled to microtubules role in membrane extrusion through tubulation under different contexts, including early phagosome tubulation, tubulation during autophagic lysosome reformation, and endosome tubulation (Du et al., 2016; Delevoye et al., 2014; Harrison et al., 2003). This likely depends on the combined action of dynein and kinesin motor proteins modulated by Rab7 and/or Arl8b GTPases (Jordens et al., 2001; Garg et al., 2011). However, a challenge in assessing the role of these proteins in phagosome resolution is that they are also required for maturation and hence, altering their function will interfere with both phases of the pathway (Harrison et al., 2003; Garg et al., 2011). The development of tools that can deactivate these proteins acutely is necessary to dissect their roles in phagosome maturation versus resolution.

Finally, by using independent compounds like ikarugamycin, pitstop, dynole 34-2, and dyngo-4a, and induced chemical displacement of clathrin, we found that clathrin and dynamin are required for budding events at the phagolysosome. The role of clathrin on lysosomes remains poorly defined, despite early observations that clathrin associates with lysosomes and that multiple adaptor protein complexes act on lysosomes or lysosome-like organelles (Arneson et al., 1999; Traub et al., 1996; Saffi and Botelho, 2019). More recently, clathrin and adaptor protein complexes like AP-2 were discovered to have a critical role in lysosome regeneration from spent autolysosomes. This requires the recruitment of distinct PIPKI isoforms to spent autolysosomes to generate a first burst of PtdIns(4,5)P_2_ that leads to assembly of clathrin to nucleate a tubule, followed by a second burst at the tips of the tubules that helps induce clathrin-mediated vesiculation of lysosome precursors (Rong et al., 2012). Whether a similar process occurs on resolving phagosomes is unknown. Ultimately, the initiation and coordination of phagosome resolution may depend on the type of phagosomal cargo, its physical parameters like particle rigidity, and catabolites generated during degradation, which may interface with mTOR, the acto-myosin cytoskeleton, and the membrane curvature machinery.

### Lysosome consumption and regeneration during phagosome resolution

We provide evidence that phagosome maturation consumes “free” lysosomes, limiting the degradative capacity of subsequently formed phagosomes. However, we also observed that this is a transient phenomenon since cells with phagosomes carrying degradable cargo quickly recovered “free” lysosomes and displayed secondary phagosomes with higher proteolytic activity compared to cells whose phagosomes had not been resolved or contained indigestible cargo.

Importantly, clathrin inhibition impaired phagosome resolution and prevented secondary phagosomes from acquiring their full degradative potential relative to control conditions. Admittedly, prolonged clathrin inhibition is a complex manipulation likely to have indirect effects. Indeed, clathrin inhibitors alone caused a reduction in the degradative capacity of phagosomes, likely by impairing trafficking of newly synthesized hydrolases (Borgne and Hoflack, 1997; Ludwig et al., 1994). This is consistent with reduced degradation of phagosomes formed after an initial round and in cells treated with cycloheximide. Nevertheless, the combined effect of clathrin inhibition and phagosomal load greatly hindered the proteolytic activity of secondary phagosomes. This suggests that failure to undergo resolution of earlier phagosomes reduced macrophage capacity to deal with subsequent phagocytosis.

Overall, we propose that phagosome resolution helps regenerate lysosomes consumed during phagosome maturation. However, we suspect that phagosome-derived vesicles are heterogeneous and may consist of distinct lysosome-like intermediates. Given that 80% of phagosome-derived fragments possessed lysosomal proteins and an acidic and proteolytic lumen, we speculate that phagosome resolution mostly regenerates endolysosome-like compartments (Bright et al., 2005). Yet, a fraction of PDVs may be akin to terminal storage lysosomes (Bright et al., 2005) or proto-lysosomes, which are depleted of degradative capacity as observed during autophagic lysosome reformation (Yu et al., 2010). Additionally, we suspect that the biosynthetic pathway and TFEB-mediated activation play a role in maintaining the degradative power of subsequent phagosomes. Thus, we propose dedicated studies to better dissect the relative contributions of phagosome resolution, lysosome reformation, and biosynthetic pathways towards maintaining macrophage degradative power after phagocytosis.

## Materials and Methods

### Cell culture, transfection and mammalian expression constructs

RAW 264.7 murine macrophages (ATCC TIB-71™, American Type Culture Collection, Manassas, VA, USA) were cultured at 37°C and 5% CO_2_ in DMEM supplemented with 10% heat-inactivated FBS (Wisent Inc., St-Bruno, QC, Canada). RAW 264.7 macrophages deleted for TFEB were a kind gift from Dr. Rosa Puertollano (NIH) and described in (Pastore et al., 2016). The clathrin light chain-GFP construct was a gift from Dr. C. Antonescu and described in (Aguet et al., 2013). The construct encoding PLCδ-PH-GFP was from Addgene and described in (Stauffer et al., 1998). pMito-mCherry-FRB (henceforth, MitoTrap) and GFP-FKBP-CLCa are plasmid constructs used for rapamycin-induced clathrin displacement (clathrin *knock-sideways*) were kindly provided by S. Royle (University of Warwick, Coventry, United Kingdom) and described (Cheeseman et al., 2013). Macrophages were transfected using FuGENE HD (Promega Corp., Madison, WI, USA) based to the manufacturer’s instructions.

### Bone marrow-derived primary macrophage culture

Bone marrow-derived macrophages (BMDMs) were harvested from wild-type 7-9-week-old female C57Bl/6 mice. Briefly, bone marrow was perfused from femurs and tibias through with phosphate-buffered saline (PBS) using a 27G syringe. Red blood cells were eliminated by osmotic lysis and BMDMs differentiated in DMEM supplemented with 10% heat-inactivated fetal bovine serum, 20 ng/mL recombinant mouse macrophage colony-stimulating factor (PeproTech, Rocky Hill, NJ), and 5% penicillin/streptomycin. Cells were incubated at 37°C with 5% CO_2_. The medium was changed every two days. Experiments were conducted on day 7 of bone-marrow isolation. All animals were used following institutional ethics requirements.

### Preparation of bacterial targets

To obtain fluorescently-labelled *E. coli*, the DH5α *E. coli* strain was transformed with the pBAD::mRFP1 plasmid (Addgene plasmid #54667; Campbell et al., 2002) or the pZsGreen vector (cat. # 632446; Takara Bio USA, Inc., Mountain View, CA, USA). Transformed *E. coli* were grown overnight at 37°C on Luria-Bertani (LB) agar plates and colonies were subsequently cultured overnight at 37°C in LB broth under agitation. The agarized and broth media were supplemented with 100 µg/mL ampicillin (Bioshop Canada Inc., Burlington, ON, Canada). For ZsGreen-*E. coli*, 1% D-glucose (Bioshop) was also added to the LB plates and broth to suppress leaky expression of ZsGreen from the *lac* operon. The overnight cultures were subcultured at 1:100 dilution and grown at 37 °C until mid-log phase, at which time mRFP1 expression was induced through supplementation with 5 mM L-arabinose (Bioshop) for 4 h, while ZsGreen expression was induced with 1 mM IPTG (Sigma Aldrich, Oakville, ON, Canada) for 3 h. *E. coli* bacteria were subsequently fixed in 4% paraformaldehyde (PFA; Electron Microscopy Sciences, Hatfield, PA, USA) in phosphate-buffered saline (PBS).

To obtain fluorescently-labelled filamentous targets, mCherry-*Lp* were prepared as described (Prashar et al., 2013). Briefly, *Lp* containing the KB288 plasmid, a gift from Dr. A.K. Brassinga and described in (Brassinga et al., 2010), were first grown at 37°C and 5% CO_2_ on buffered charcoal-yeast extract plates for 3 days. Colonies were grown for an additional 24 h at 37°C in buffered yeast extract (BYE) media under agitation. The *Lp* filaments were then killed with 4% PFA solution. For the analysis of phagosome resolution utilizing filaments, bacteria longer than 15 µm were considered filamentous (Prashar et al., 2013).

### Phagocytosis assays

For the phagocytosis of *Lp* filaments, the PFA-killed bacteria were opsonized with 0.1 mg/mL of anti-*Legionella* antibody (Public Health Ontario, Toronto, ON, Canada) or 4 mg/mL for human IgG for 1 h at room temperature (RT; Prashar et al., 2013). RAW macrophages were pre-cooled to 15°C for 5 minutes before the cells were challenged with filaments. Bacterial attachment was synchronized by spinning the cells at 300 xg for 5 minutes at 15°C. Following a 15-minute incubation at 37°C, unbound filaments were washed off with PBS. Macrophages were subsequently incubated at 37°C to allow phagocytosis to progress to the indicated time points, at which time the cells were either fixed with 4% PFA for 20 minutes or imaged live.

For the phagocytosis of *E. coli* rods and beads, the bacteria, 3.87 µm polystyrene latex beads (Bangs Laboratories, Fishers, IN, USA) or 3.0 µm polystyrene latex beads (Sigma Aldrich) were opsonized with 4 mg/mL of human IgG (Sigma Aldrich) for 1 h at RT. To coat beads with DQ-BSA, a 5% w/v suspension of 3.0 µm beads or 0.5% w/v suspension of 2.0 µm Dragon Green fluorescent beads (Bangs Laboratories) was incubated with 0.5 mg/mL DQ-Red BSA (ThermoFisher Scientific, Mississauga, ON, Canada) for 1 h before opsonization with human IgG or 200 µg/mL anti-BSA antibody (Invitrogen, Burlington, ON, Canada). RAW cells were pre-cooled to 4°C for 5 min prior to the cells being presented with targets at ratios of 20-400 rods per cell, 10 3.87 µm beads per cell or 25-135 3.0 µm beads per cell, depending on if phagosomal saturation was required. Target attachment was synchronized by spinning cells at 300 xg for 5 min at 4°C. Following a 15- to 60-minute incubation at 37°C, unbound targets were washed off with PBS. The cells were then incubated at 37°C to allow phagocytosis to progress to the indicated time points and either fixed with PFA or imaged live. For assays using two-rounds of phagocytosis, the procedure for both the first and second rounds of phagocytosis was the same as described above, except the secondary phagocytic challenge was done after the first round of phagocytosis at specified times. The secondary challenge was then followed with no chase or 1 h chase to elicit further maturation, prior to fixation or live cell imaging.

For the phagocytosis of bead clumps, 0.1 µm Tetraspeck microspheres (blue/green/orange/dark red; Thermo Fisher Scientific) were opsonized with 4 mg/mL of human IgG for 1 hour at RT before leaving the mixture at 4°C overnight. Macrophages were pre-cooled to 4°C for 5 min prior to the cells being presented with bead clumps at ratios of 1,400 beads per cell. Bead clumps were spun onto the cells at 300 x*g* for 5 min at 4°C. Following a 15-minute incubation at 37°C, unbound bead clumps were washed off with PBS. Macrophages were subsequently incubated at 37°C to allow phagocytosis to progress to the indicated time points, at which point the cells were imaged live.

### Exocytosis of phagosomal content

Following phagocytosis of mRFP1-labeled *E. coli* and incubation for 1 h, 6 h, and 24 h, the cell media was centrifuged at 10,000 xg for 3 min to remove loose bacteria and cellular debris. To assess mRFP1 in the cell-free supernatant, proteins were precipitated using ice-cold trichloroacetic acid (Bioshop) added to a final concentration of 10%. The mixture was incubated on ice for 20 min at RT and centrifuged at 17,000 xg for 2 min. The TCA precipitate was washed twice with ice-cold acetone (Bioshop), resuspended by sonication and centrifuged at 17,000 xg for 2 min. Pellets were dried and then resuspended in 1x Laemmli.

To asses mRFP1 in phagosomes, cells were washed 3x with ice-cold PBS treated to remove bacteria adhered to the cells. Cells were then washed 3x with ice-cold PBS and lysed with 1x Laemmli. 50 µL of mRFP1-labeled *E. coli* at 1 OD was also lysed with 1x Laemmli.

The contents of the *E. coli*, sample media, and cells were analyzed by Western Blot, using the rabbit anti-RFP antibody (1:1,000, Rockland Immunochemicals, Inc., Gilbertsville, PA, USA) and GAPDH using the rabbit anti-GAPDH antibody (1:1,000, Cell Signalling Technologies). HRP-conjugated anti-rabbit antibody (Bethyl) was used at 1:10,000 and developed with Clarity Western ECL Substrate (Bio-Rad). mRFP1 from phagolysosomes was identified by the increased gel migration compared to native mRFP1 (Katayama et al., 2008).

### Isolation of phagosomes

To isolate phagosomes, a sucrose gradient was prepared by adding 1 mL of 60% sucrose suspension to a 1 mL ultracentrifuge tube and the solution was centrifuged at 50,000 xg for 1 h at 4°C. Following centrifugation, the sucrose gradient was carefully placed on ice until use.

Macrophages were allowed to internalize IgG-opsonized polystyrene beads for 60 min. Following phagocytosis, the cell media was replaced with cold homogenization buffer (20 mM Tris (Bioshop), 1:400 protease inhibitor cocktail (Sigma), 1 mM AEBSF (Bioshop), 1 mM MgCl_2_ (Thermo Scientific), 1 mM CaCl_2_ (Biobasic, Markham, ON, Canada), 0.02 mg/mL RNase (Roche Diagnostics, Laval, QC, Canada), and 0.02 mg/mL DNase (Roche), pH 7.4), and the cells were dislodged by mechanical scraping. The cell suspension was centrifuged at 500 xg for 5 minutes at 4°C, and the pellet was resuspended in homogenization buffer. Cells were lysed by passing the suspension through a 22-gauge needle 5-10 times, then the lysate was centrifuged at 1000 xg for 5 min. The pellet was resuspended in 4% PFA and incubated at room temperature for 20 min to fix the lysate and secure protein complexes on the phagosome surface. The pellet was washed 3 times with PBS followed by centrifugation at 3000 xg for 5 min per wash, then resuspended in 200 µL of PBS. To separate phagosomes from the resuspended lysate pellet, the resuspended pellet was loaded onto the sucrose gradient and then centrifuged at 21,000 xg for 10 min at 4°C. The bead-containing fraction was extracted from the gradient with a 22-gauge needle and washed with ice-cold PBS.

### Pharmacological inhibitors

Inhibitors and the DMSO (Bioshop) vehicle control were applied post-phagocytosis and maintained until fixation or the conclusion of the experiment. To increase cytosolic Ca^2+^, cells were incubated with 10 µM of ionomycin (Sigma-Aldrich) and 1.2 mM of CaCl_2_ (Bioshop) 1-2 h post-phagocytosis for up to 1 h. For pH neutralization of the phagolysosome, macrophages were treated with 1 µM concanamycin A (ConA; Bioshop) and 10 mM NH_4_Cl (Bioshop) 40 min after the start of phagocytosis. In order to inhibit proteases, cells were incubated with a protease inhibitor cocktail (Sigma-Aldrich) 40 min after phagocytosis, which included 1.0 mM AEBSF, 0.8 µM aprotinin, 40.0 µM bestatin, 14.0 µM E-64, 20.0 µM leupeptin and 15.0 µM pepstatin A. To inhibit the cytoskeleton and dynamin, macrophages were treated with the following inhibitors 40 min post-phagocytosis: 10 µM nocodazole (Sigma-Aldrich), 10 µM taxol (Sigma-Aldrich), 2 µM cytochalasin D (EMD Millipore, Oakville, ON, Canada), 1 µM jasplakinolide (EMD Millipore), 5 µM dyngo-4a (Abcam) and 5 µM dynole 34-2 (Abcam). For clathrin inhibitors, cells were incubated with 10 µM of pitstop 2 (Abcam, Cambridge, MA, USA) or 0.5-2.0 µg/mL ikarugamycin (Sigma-Aldrich) 40 min to 4 h after the start of phagocytosis. To inhibit *de novo* protein synthesis, macrophages were treated with 1 µM cycloheximide (Bioshop) 1h after phagocytosis.

### Rapamycin-induced clathrin displacement (clathrin knock-sideways)

After 24 h of transfection, cells were treated with 1 μM rapamycin or DMSO for 1h. After 1h, 0.5 OD of non-labeled *E. coli* were added into each well +/- rapamycin and pulsed for 30 min at 37 ^0^C and 5% CO_2_. Cells were washed 3x with 1x PBS and warm medium was added and chased for 3 h. For staining internalized *E. coli,* PFA-fixed cells were permeabilized with 0.1% Triton for 20 min at RT. Cells were washed 3x with 1x PBS and incubated at RT with 1:100 anti-*E. coli* antibody prepared with 1% BSA. Cells were washed with 1x PBS for 3x, and incubated with 1:1000 fluorescent secondary antibody in 1% BSA for 1h at RT. After the last wash, coverslips were mounted using Dako mounting medium and imaged. Number of fragments in control and rapamycin treated cells were counted using ImageJ.

### Immunofluorescence and fluorescence labeling

External beads were stained for live-cell imaging by incubating macrophages with Cy2-conjugated anti-human IgG (1:100; Jackson ImmunoResearch Labs, West Grove, PA, USA) for 30 min on ice. For internal staining, PFA-fixed cells were permeabilized with 0.1% Triton X-100 for 20 min with the exception of staining with the LAMP1 antibody, which required permeabilization with ice cold methanol for 5 min instead. Primary antibodies that were diluted in 5% skim milk or 1% BSA solutions were applied for 1 h at RT and included: anti-*Legionella* antibody (1:3000; Public Health Ontario), anti-*E. coli* antibody (1:100, Bio-Rad), anti-LAMP1 (1:200; 1D4B; Developmental Studies Hybridoma Bank, Iowa City, IA) and anti-LAMP2 antibody (1:100; ABL-93; Developmental Studies Hybridoma Bank, Iowa City, IA, USA).

Fluorescent secondary antibodies (ThermoFisher Scientific or Bethyl Laboratories, Montgomery, TX, USA) were utilized at a 1:1000 dilution within 5% skim milk or 1% BSA solutions for 1 h at RT. The coverslips were mounted using Dako fluorescence mounting medium (Agilent Technologies, Inc. Mississauga, ON, Canada).

For labelling acidic compartments, cells were stained with Lysotracker Red DND-99 at 1 µM for 1 h (ThermoFisher Scientific). To label degradative compartments, cells were labelled with Magic Red Cathepsin-L substrate (ImmunoChemistry Technologies, LLC, Bloomington, MN, USA) for 1 h, as per the manufacturer’s instructions.

For labeling isolated phagosomes, phagosomes were incubated with the following primary antibodies: anti-LAMP1 (1:200 in PBS) and anti-clathrin heavy chain (1:200 in PBS; D3C6; Cell Signalling Technology, Danvers, MA, USA). Phagosomes were washed 3 times with 0.5 % BSA followed by centrifugation at 2000 xg for 1 min. Fluorescent secondary antibodies were added to the phagosomes at a 1:1000 dilution and incubated for 1 h at room temperature, then phagosomes were washed 3 times with 0.5 % BSA followed by centrifugation at 2000 xg for 1 min. Phagosomes were resuspended in PBS and mounted with Dako mounting medium.

### Microscopy

Confocal images were acquired using two different spinning disc confocal microscope systems. The first was a Quorum Wave FX-X1 spinning disc confocal microscope system consisting of an inverted fluorescence microscope (DMI6000B; Leica Microsystems, Concord, ON, Canada) equipped with a Hamamatsu ORCA-R^2^ camera and a Hamamatsu EM-CCD camera (Quorum Technologies, Inc., Guelph, ON, Canada). The second was a Quorum Diskovery spinning disc confocal microscope system consisting of an inverted fluorescence microscope (DMi8; Leica) equipped with the Andor Zyla 4.2 Megapixel sCMOS and Andor iXON 897 EMCCD camera (Quorum Technologies, Inc.). We also used an inverted microscope (IX81; Olympus Life Science, Richmond Hill, ON, Canada) equipped with a Rolera-MGI Plus EM-CCD camera (Q Imaging, Tucson, AZ, USA). The microscope systems were controlled by the MetaMorph acquisition software (Molecular Devices, LLC, San Jose, CA, USA). Images were acquired using a 63x oil immersion objective (1.4 NA), a 40x oil immersion objective (1.3 NA) or a 40x dry objective (0.60 NA). For live-cell imaging, coverslips were imaged in a chamber containing DMEM supplemented with 10% FBS in a microscope-mounted chamber maintained at 37 °C and 5% CO_2_.

### Live-cell imaging of clathrin-GFP and phagosome resolution

One day post-transfection, RAW cells were treated with 1 μM rapamycin or DMSO for 1 h and challenged of *E. coli* bacteria to trigger phagocytosis. After 30 min, the bacteria in the media was removed by washing the cells 3x with PBS and warm culture media was added to allow phagocytosis maturation for 1-2 h period. Live-cell imaging was performed, keeping cells at 5% CO_2_ and 37 °C in an environmental control chamber with a Quorum Diskovery spinning disc confocal microscope set to acquire 5-plane z-stack at 0.4 µm interval at 1 frame/5 s for 5 min periods. Imaging analysis was done with ImageJ (NIH, Bethesda, MD) by counting the number of clathrin-positive and clathrin-negative fission events (tubule formation, budding, constriction).

### Image processing and analysis

Image processing and quantitative analysis were performed using FIJI (Schindelin et al., 2012) or Volocity (PerkinElmer, Waltham, MA, USA), where image enhancements were completed without altering the quantitative relationship between image elements. For the quantification of the total volume of *E. coli*-containing phagosomes in each cell, phagosomes were first identified in Volocity by their high fluorescence intensity and then by their volume, set as greater than 1 μm^3^. The volume of phagosomes in each cell were added together for each time point and normalized to the first acquisition time point (T_0_). For the quantification of the total volume of PDVs containing *E. coli*-derived debris, we had Volocity select dimmer fluorescence objects and then defined their volume as greater than 0.02 μm^3^, but less than 5 μm^3^. The volume of individual PDVs in each cell were summed and in the case of time-lapse microscopy, normalized to the first time point. Phagosomes were excluded from the PDV quantifications, as the inclusion of lower fluorescence intensities allowed PDVs closely surrounding the phagosome to be included in the phagosome volume, thereby increasing the apparent size of the phagosomes above the volume threshold for PDVs (Supplemental Figure S5). For the quantification of the total volume of *Lp*-containing phagosomes and PDVs in each cell, we utilized the same methodology as for *E. coli*, except that the phagosome was included in the quantifications if its volume exceeded 5 µm^3^. For the co-occurrence of Lysotracker red or Magic red with PDVs containing *E. coli*-derived debris, we used Manders’ colocalization analysis in Volocity, where PDVs were included within this analysis if their surface area was greater than 0.1 µm^2^, but less than 4 µm^2^. The markers were considered to co-occur when the M_2_ colocalization coefficient was determined to be greater than 0.7. We then reported the percentage of PDVs that were positive for Magic red or Lysotracker (co-occurrence of markers).

For the counting of phagosome and PDV events, images of mRFP1-labeled *E. coli* and PDV were first thresholded to exclude background and macrophage autofluorescence signals, and to minimize signal overlap of events while retaining as much low-intensity events as possible. Once an intensity threshold was identified, it was applied to all images of all samples of the same experimental replicate. External events were identified by colocalization with external anti-*E. coli* fluorescence signal and excluded from analysis. The thresholded images were used to create binary images of the samples, and the binary images were processed by object segmentation using FIJI’s Watershed function. Intact phagosome events were identified manually by the rod-shaped events characteristic of *E. coli* shape. PDVs between 0.015 µm^2^ and 1.2 µm^2^ were counted using FIJI’s Analyze Particles function.

For the quantification of the number of free lysosomes in each cell, we used Volocity to separate touching objects in the LAMP1 channel using an object size guide of 0.29 µm^3^ (determined by assessing the lysosome size in resting macrophages). LAMP1-positive objects were considered free lysosomes if their volume was greater than 0.02 µm^3^, but less than 5 µm^3^ and they were not touching the filament-containing phagolysosomes (applied a mask to the filament). The treatments were finally normalized to the “no-phagocytosis” group, which was considered to contain 100% free lysosomes.

For the determination of presence and absence of LAMP1 and LAMP2 on PDVs, first, PDVs in an image were identified as vesicles with an image area between 0.015 µm^2^ and 1.2 µm^2^. Then PDVs were assessed for the presence of absence of LAMP1 or LAMP2 colocalized to the PDV, and cells are scored by percentage of PDVs with colocalized LAMP1 or LAMP2 fluorescent signal. For the determination of phagosome mixing, images of second-wave phagosomes and fragments (GFP) were thresholded to exclude background and macrophage autofluorescence signals, then a binary mask was formed.

Second-wave events external to the cell were determined by comparison to Brightfield images and excluded from the mask. The mean fluorescence intensity of first-wave phagosome remnants (mRFP1) colocalized to the second-wave mask was determined and corrected by numerical subtraction of mean background intensity. For the determination of DQ-BSA intensity, images in a stack corresponding to the lower half of DQ-BSA beads were collapsed by Maximum Intensity Projection (MIP), and the MIP images were thresholded to exclude non-DQ-BSA bead events. External beads were identified by colocalization of Cy2-anti-human or Cy-3-anti-human fluorescence signal and excluded from analysis by relevant mask subtraction.

Following mask subtraction, DQ-BSA proteolytic activity was determined by measuring the mean fluorescence of internalized beads and corrected by numerical subtraction of mean external DQ-BSA bead intensity. Since there are multiple sources of heterogeneity (uptake of primary particles, age and resolution of phagosomes, uptake of secondary phagosomes, age of secondary phagosomes, and level of DQ-BSA labelling of beads), we analysed the entire sampled population rather than sampled mean. Deconvolution (30 iterations) was performed utilizing Volocity software. Figures were assembled using Adobe Illustrator (Adobe Systems, Inc., San Jose, CA, USA).

### Statistical analysis

Unless otherwise indicated, data are presented as mean ± SEM of at least three independent experiments unless otherwise stated. The numbers of cells assessed in each experiment is indicated within the figure legends. Statistical analysis was performed using GraphPad Prism software (GraphPad Software, Inc., San Diego, CA, USA). Unless otherwise indicated, an unpaired, two-tailed Student’s t-test was utilized to compare two conditions, while multiple conditions were compared using a one-way ANOVA with Tukey’s *post hoc* test, a two-way ANOVA with Tukey’s *post hoc* test or a two-way ANOVA with Sidak’s *post hoc* test. Curves were fit to the data using non-linear regression, where AIC was used to select the model that best fit the data. The slopes of the lines using linear regression were statistically compared using an ANCOVA. In any of the statistical tests performed, values of p < 0.05 were considered significant.

## Abbreviations

Arl 8: Arf-like GTPase 8

CLC-GFP: GFP-fusion of the clathrin-light chain

ConA: concanamycin A

DQ-BSA: dye-quenched bovine serum albumin

IKA: ikarugamycin

LAMP1: lysosomal-associated membrane protein 1

LAMP2: lysosomal-associated membrane protein 2

LB: Luria-Bertani medium

Lp: *Legionella pneumophila*

mTORC1: mechanistic target of rapamycin complex 1

PDV: phagosome-derived vesicle

PtdIns(3)P: phosphatidylinositol 3-phosphate

PtdIns(3,5)P2: phosphatidylinositol 3,5-biphosphate

PtdIns(4)P: phosphatidylinositol 4-phosphate

TFEB: transcription factor EB

## Supplemental Materials

**Supplemental Figure S1: Phagolysosomes do not undergo exocytosis, but fragment. A.** One-hour post-phagocytosis of IgG-opsonized beads RAW cells were treated with 10 µM ionomycin, in the presence of 1.2 mM of Ca^2.^and live cell DIC images were acquired at the indicated time points. Scale bars are 10 µm. **B.** Number of latex beads per cell at the indicated treatment times, normalized to the number of beads/cells observed at the onset of the treatment (T_0_). Data are mean ± SEM from groups of 10 cells measured in 3 independent experiments. See video 2 for corresponding movies. **C**. RAW cells expressing PLCδ1-PH-GFP showing IgG-opsonized beads in phagosomes, treated with 10 µM ionomycin or DMSO (control) in the presence of 1.2 mM of Ca^2+^. Images were acquired before or after 10 minutes of treatment. Asterisks correspond show the position of the beads shown in the insets. Scale bars are10 µm. **D**. Live cell imaging time series of IgG-aggregated fluorescent 0.1 µm latex beads after 2 h phagocytosis by RAW cells. Beads are depicted in a rainbow scale. Red and blue correspond to the highest and lowest florescence intensity level, respectively. The smaller panels show the far-red and DIC channels for the first frame of the movie. The cell contour is delineated with white dots. Scale bar: 5 µm. Stills are from Video 5. **E**. Macrophages were assessed for exocytosis of digested phagosomal contents. RAW macrophages internalized mRFP1-labeled *E. coli* for 1 h, then incubated for the indicated timepoints. TCA-precipitates of cell media and cell lysates were probed for mRFP1, detecting a native and a cleaved mRFP1 product that accumulated overtime. GAPDH was used as a loading control and to detect cell lysis. **F.** Quantification of cleaved-mRFP1 levels normalized to GAPDH in media and within cells. Data shown as mean ± STD of n=6 independent experiments. Conditions were compared using one-way ANOVA and Holm-Sidak’s post-hoc test. *: p<0.05, ns: not significant.

**Supplemental Figure S2: Clathrin inhibition blocks phagosome resolution. A.** Single-cell live cell imaging time series of RAW cell expressing clathrin light chain-GFP showing internalized IgG-opsonized beads at 3 h post-phagocytosis. Colour scale indicates the fluorescence intensity of clathrin-GFP. Time interval between frames is 10 s. * indicates internal beads and green arrows indicate accumulation of clathrin around phagosomes. Scale bar is 5 µm. **B.** RAW cells internalized mRFP1-*E. coli* for 1 h before treatment with Ikaguramycin. Live-cell imaging commenced after uptake (0 h) or >6 h after phagocytosis. Scale bar: 10 µm. **C.** Intact phagosomes and PDVs per cell were quantified as puncta (particle number) for experiments displayed in B and normalized to 0 h. Data shown as mean ± SEM of 15 images per control/treatment across 3 independent experiments, where each image display 8 to 29 cells. Control and inhibitor-treated cells at each time point were compared statistically using a two-way ANOVA with Sidak’s post-hoc test (*, p < 0.05; ns, not significant). **D.** RAW macrophages internalized mRFP1-*E. coli*, chased for 1 h, and then treated with either vehicle or 10 µM Pitstop 4 or 6 h post-phagocytosis. **F.** Primary macrophages were presented with mRFP1-*E. coli* and were treated either with vehicle, Pitstop, or ikarugamycin post-maturation. Cells were fixed and imaged 3h post-phagocytosis. **E** and **G**. PDVs/cell were quantified in 60 cells per condition across three independent trials and compared statistically using a one-way ANOVA with Dunnett’s multiple comparison test. (*, p<0.001; **, p<0.0001).

**Supplemental Figure S3. A** and **C.** 40 min post-phagocytosis of mRFP1-*E. coli*, RAW cells were treated with the dynamin inhibitors, dyngo-4a, dynole 34-2, or vehicle (DMSO), fixed at the indicated time points and immunostained for *E. coli. E. coli* fluorescence labelling is shown in rainbow scale. Red and blue are the highest and lowest intensity level, respectively. Dotted lines indicate the boundary of the cells. Scale bar: 5 μm. **B** and **D.** The total volume of PDVs per cell for indicated treatments. Data are means ± SEM of sample of 25 cells from 3 independent experiments. Treatments were compared statistically using a one-way ANOVA with Tukey’s post-hoc test. Conditions with different letters indicate statistically different (p < 0.05). **E.** Macrophages treated with dynole 34-2 or DMSO (vehicle) internalized Alexa Fluor 546-labeled transferrin for 10 minutes prior to fixation. White dots indicate cell boundaries. Scale bar: 30 μm.

**Supplemental Figure S4. The role of the biosynthetic pathway and TFEB in phagosome resolution and degradative capacity of macrophages.** A. RAW cells internalized mRFP1-*E. coli* for 20 minutes, and after a 1h chase, cells were treated with vehicle or cycloheximide (CHX) for 6 h total. B. Quantification of PDVs per cell from A show as means ± SEM from groups of 15 images, each containing between 5 to 15 cells, from 3 independent experiments. Statistical analysis by one-way ANOVA with Tukey’s post-hoc test (*, p < 0.05; **, p< 0.01; ***, p<0.001; ns, not significant). C. Wild-type and TFEB-deleted macrophages were either given no bacteria or allowed to internalize unlabelled *E. coli* for 1 and 6 h as a first-round of phagocytosis prior to the addition of DQ-BSA-opsonized beads (red) for a second-round of engulfment. Live-cell imaging was performed after immunostaining of external beads (green). The lower panel shows the DQ-BSA in fire scale. Scale bar: 10 µm. D. Quantification of DQ-BSA fluorescence intensity of 120 cells per condition across three independent experiments. Fluorescence intensity of internalized DQ-BSA was measured and subtracted from the mean intensity of external DQ-BSA beads, and the difference between the conditions were statistically tested using a two-way ANOVA and Tukey’s post-hoc test (***, P< 0.0001; ns, not significant).

**Supplemental Figure S5: Overview of the quantification method used to determine the total volume of phagosomes and PDVs in each cell. A.** A still image from the same video displayed in Figure 1D is used to exemplify the quantification method. The main panel shows the red channel in a rainbow scale, where red is the highest intensity level and blue is the lowest intensity level. The smaller panels show the red and DIC channels. The white dotted line indicates the boundary of the cell and the white boxes indicate the positions of the insets. Arrows within the insets point to phagosomes, while arrowheads point to PDVs. Scale bars: main panels: 10 μm; insets: 2 μm. **B.** To quantify the total volume of phagosomes in each cell, we drew a region of interest around the cell and instructed Volocity to select objects within that region, which had a high fluorescence intensity and a volume greater than 1 μm^3^ (phagosomes are under the masked areas). The volume of phagosomes in each region of interest were summed to determine the total volume of phagosomes in each cell. The volumes of the phagosomes displayed in the insets is indicated. **C.** To quantify the total volume of PDVs in each cell, we instructed Volocity to select dimmer objects within the region of interest enclosing the cell, which had a volume greater than 0.02 μm^3^, but less than 5 μm^3^ (PDVs are under the masked areas). The volume of PDVs in each region of interest were summed to determine the total volume of PDVs in each cell. Phagosomes were excluded from the PDV quantification, since the inclusion of lower fluorescence intensity PDVs immediately surrounding the phagosomes were included in the phagosomal volume, which increased the apparent size of the phagosomes above the PDV threshold. **D.** To demonstrate that the enlarged phagosomes are excluded from the PDV quantification (in C.), we modified the PDV settings to select the objects that had a volume greater than 5 μm^3^. The volumes of these enlarged phagosomes are indicated for comparison with phagosomes quantified in B.

**Supplemental Video 1. Bead-containing phagosomes remain within macrophages over 24 h.** RAW macrophages were allowed to internalize IgG-opsonized latex beads for 2 h before cells were moved to a pre-warmed (37°C) microscope stage. Images were acquired at a rate of 1 frame/h for a 24-hour period. Still frames from Video 1 are displayed in Figure 1A. Scale bar: 10 µm.

**Supplemental Video 2. Ionomycin treatment does not trigger exocytosis of bead-containing phagosomes.** IgG-opsonized latex beads were presented to RAW cells for 1 h before cells were moved to a pre-warmed (37°C) microscope stage. After acclimatization, the cells were exposed to media containing 10 µM of ionomycin and 1.2 mM of Ca^2+^ and imaging commenced immediately. Images were acquired at a rate of 12 frames/h for a 1-hour period. Still frames from Video 2 are displayed in Supplemental Figure S1A. Scale bar: 2 µm.

**Supplemental Video 3. Phagosomes containing mCherry-*Legionella* undergo fragmentation.** RAW macrophages were challenged with IgG-opsonized, filamentous mCherry-*Legionella* for 7 h before the cells were moved to a pre-warmed (37°C) microscope stage.

Images were acquired at a rate of 6 frames/h for a 13-hour period. The mCherry fluorescence intensity is displayed as a rainbow scale, where red is the highest intensity level and blue is the lowest intensity level. Still frames from Video 3 are displayed in Figure 1C. Scale bar: 2 µm.

**Supplemental Video 4. Fission of phagosomes containing ZsGreen-*E. coli*.** Macrophages were allowed to internalize ZsGreen-*E. coli* and imaging was started immediately. Images were acquired at a rate of 2 frames/h for a 10-hour period. Scale bar = 10 µm. Video 4 is complementary to Figure 1D. Scale bar: 2 µm.

**Supplemental Video 5. Phagosomes containing fluorescent bead clumps undergo fragmentation.** Cy5-labelled 0.1 µm beads were clumped using human IgG and were presented to RAW macrophages for 2 h before the cells were moved to a pre-warmed (37°C) microscope stage. Images were acquired at a rate of 3 frames/h for a 6-hour period. The Cy5 fluorescence intensity is illustrated as a rainbow scale where red is the highest intensity level and blue is the lowest intensity level. Still frames from Video 5 are displayed in Supplemental Figure S1D. Scale bar: 2 µm.

**Supplemental Video 6. Budding of phagosomes containing intact mRFP1-*E. coli*.** Macrophages were allowed to internalize mRFP1-*E. coli* and imaging was started 15 min after phagocytosis. mRFP1-positive vesicles presented as pseudocolour to enhance visibility of low-intensity events. Images were acquired at 1 frame every 3 seconds. Still frames from Video 6 are displayed in Figure 3A. Scale bar: 2 µm.

**Supplemental Video 7. Budding of phagosomes containing degraded mRFP1-*E. coli.*** Macrophages were allowed to internalize mRFP1-*E. coli* and imaging was started 180 minutes after phagocytosis. mRFP1-positive vesicles presented as pseudocolour to enhance visibility of low-intensity events. Images were acquired at 1 frame every 3 seconds. Still frames from Video 7 are displayed in Figure 3A’. Scale bar = 2 µm.

**Supplemental Video 8. Tubulation of phagosomes containing mRFP1-*E. coli.*** Macrophages were allowed to internalize mRFP1-*E. coli* and imaging was started 180 minutes after phagocytosis. mRFP1-positive vesicles presented as pseudocolour to enhance visibility of low-intensity events. Images were acquired at 1 frame every 3 seconds. Still frames from Video 7 are displayed in Figure 3B. Scale bar: 2 µm.

**Supplemental Video 9. Splitting of phagosomes containing mRFP1-*E. coli.*** Macrophages were allowed to internalize mRFP1-*E. coli* and imaging was started 150 minutes after phagocytosis. mRFP1-positive vesicles presented as pseudocolour to enhance visibility of low-intensity events. Images were acquired at 1 frame every 3 seconds. Still frames from Video 9 are displayed in Figure 3C. Scale bar: 2 µm.

**Supplemental Video 10: Clathrin-GFP puncta associates with phagosomes and phagosome fission sites.** RAW macrophages expressing clathrin light chain-GFP engulfed mRFP1-*E. coli*. After at least 1 h of phagosome maturation, cells were imaged by spinning disc confocal acquiring 5 z-planes spaced at 0.4 µm every 5 s for 5 min. Shown are collapsed z-stacks of the green and red channels. Still frames from Video 10 are displayed in Figure 4B.

**Supplemental Video 11. Inhibition of clathrin using pitstop prevents the fragmentation of phagolysosomes containing mCherry-*Legionella*.** RAW macrophages were allowed to internalize IgG-opsonized, filamentous mCherry-*Legionella* for 4 h before cells were moved to a pre-warmed (37°C) microscope stage. After acclimatization, the cells were exposed to media containing vehicle control (DMSO) or 10 µM of pitstop 2 and imaging started 10 min after treatment. Images were acquired at a rate of 12 frames/h for a 9.5-hour period. The mCherry fluorescence intensity is illustrated in rainbow scale, where red is the highest intensity level and blue is the lowest intensity level. Still frames from Video 10 are displayed in Figure 5A. Scale bar: 2 µm.

