## Supplementary figures and images for "Phagosome resolution regenerates lysosomes and maintains the degradative capacity in phagocytes"

### Supplemenatl Figure S1

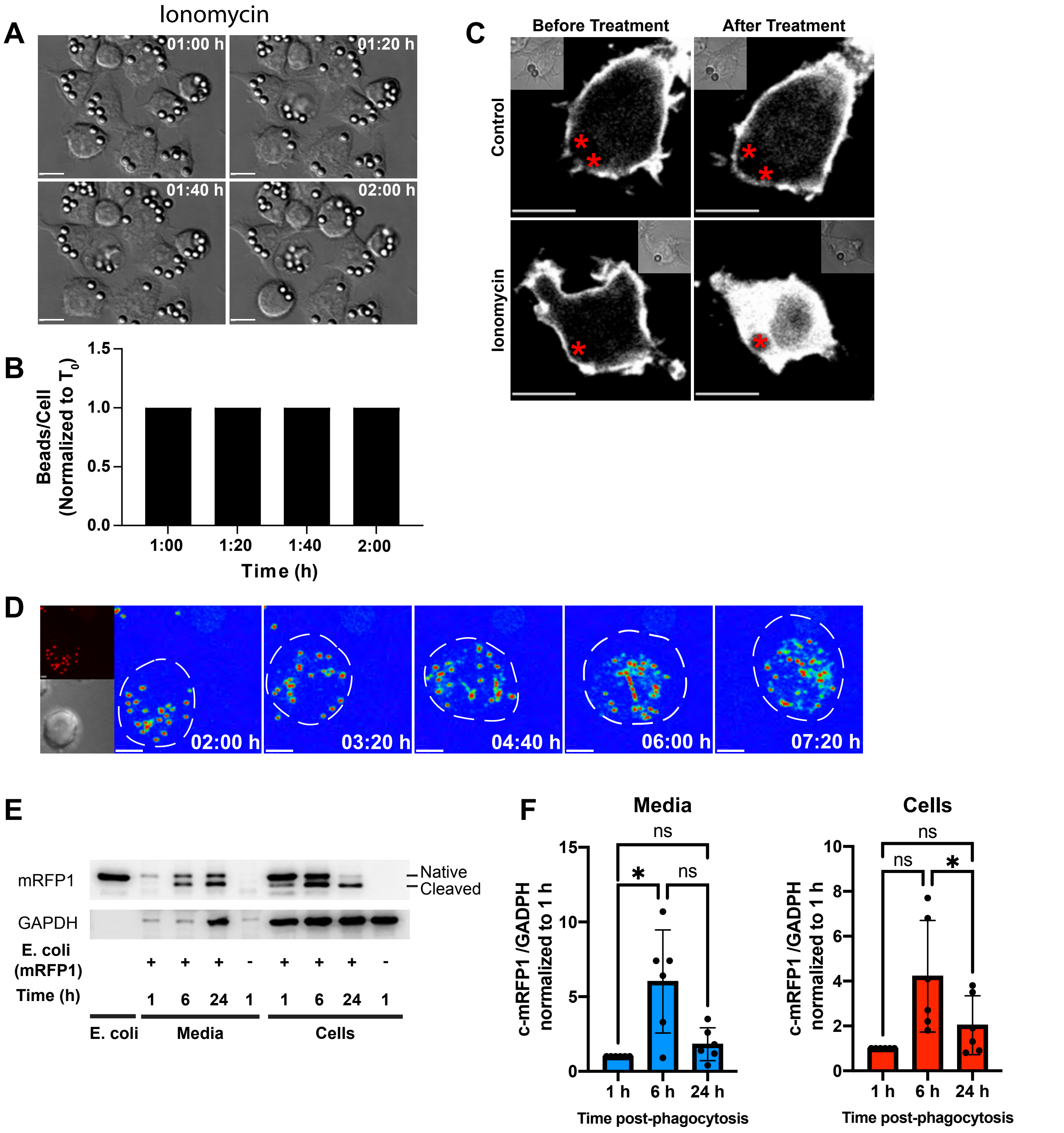


Supplemental Figure S1

### Supplemenatl Figure S2

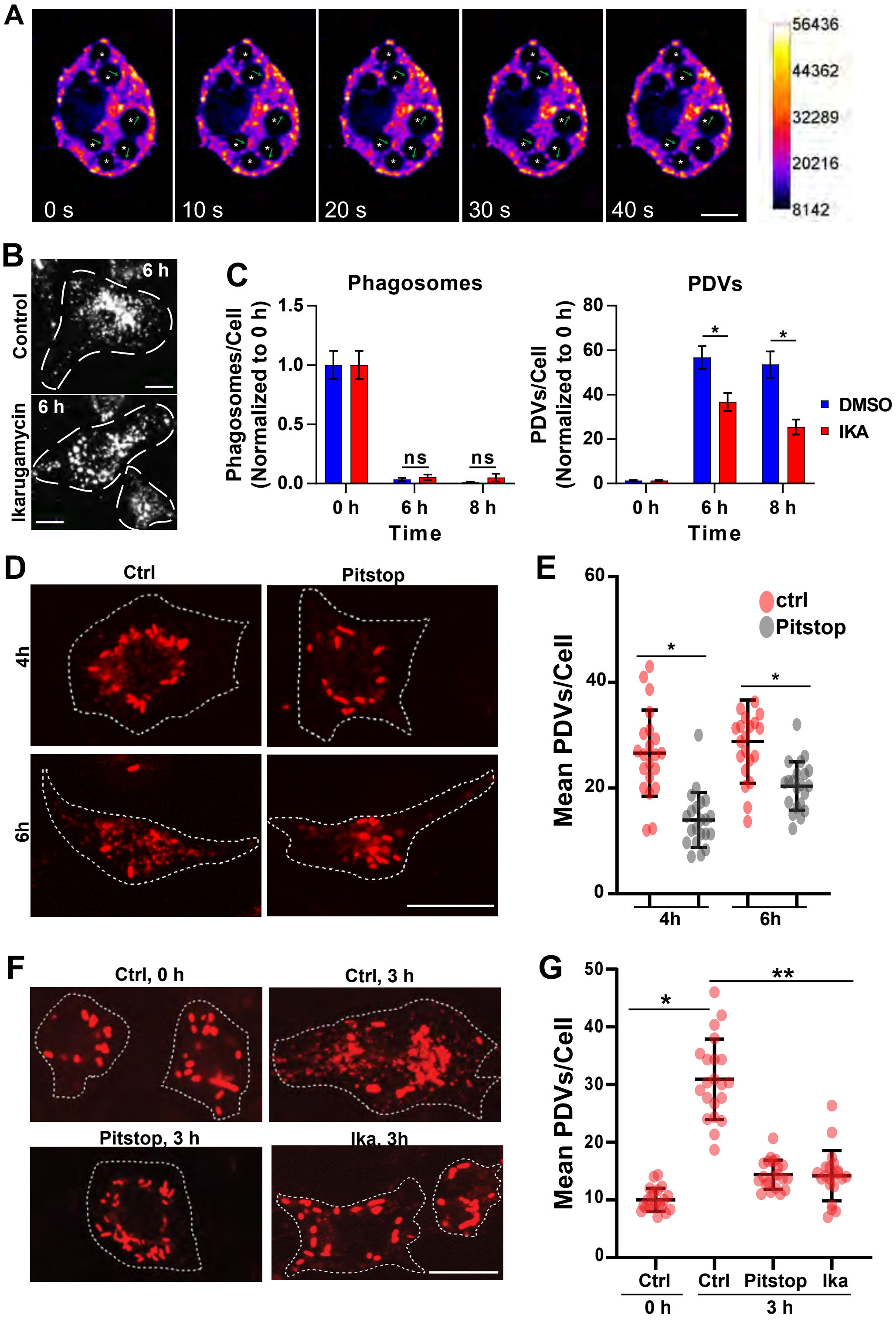


Supplemental Figure S2

### Supplemenatl Figure S3

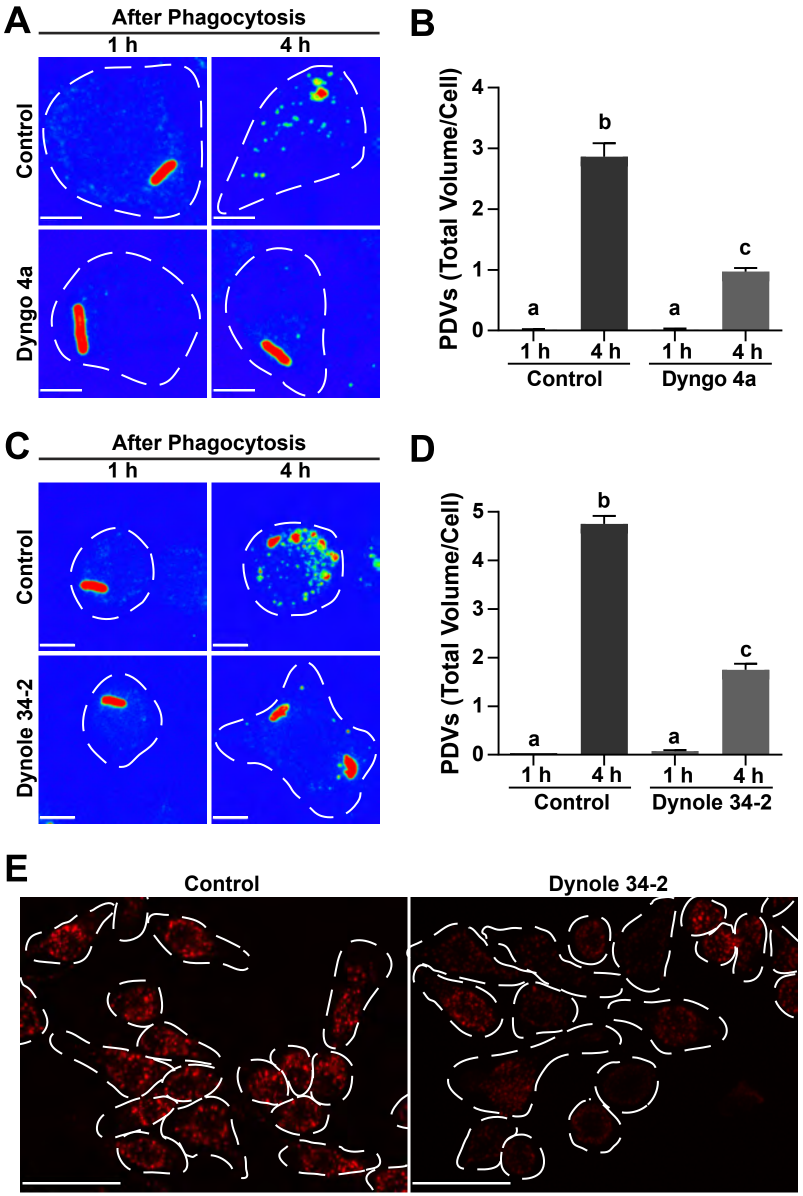


Supplemental Figure S3

### Supplemenatl Figure S4

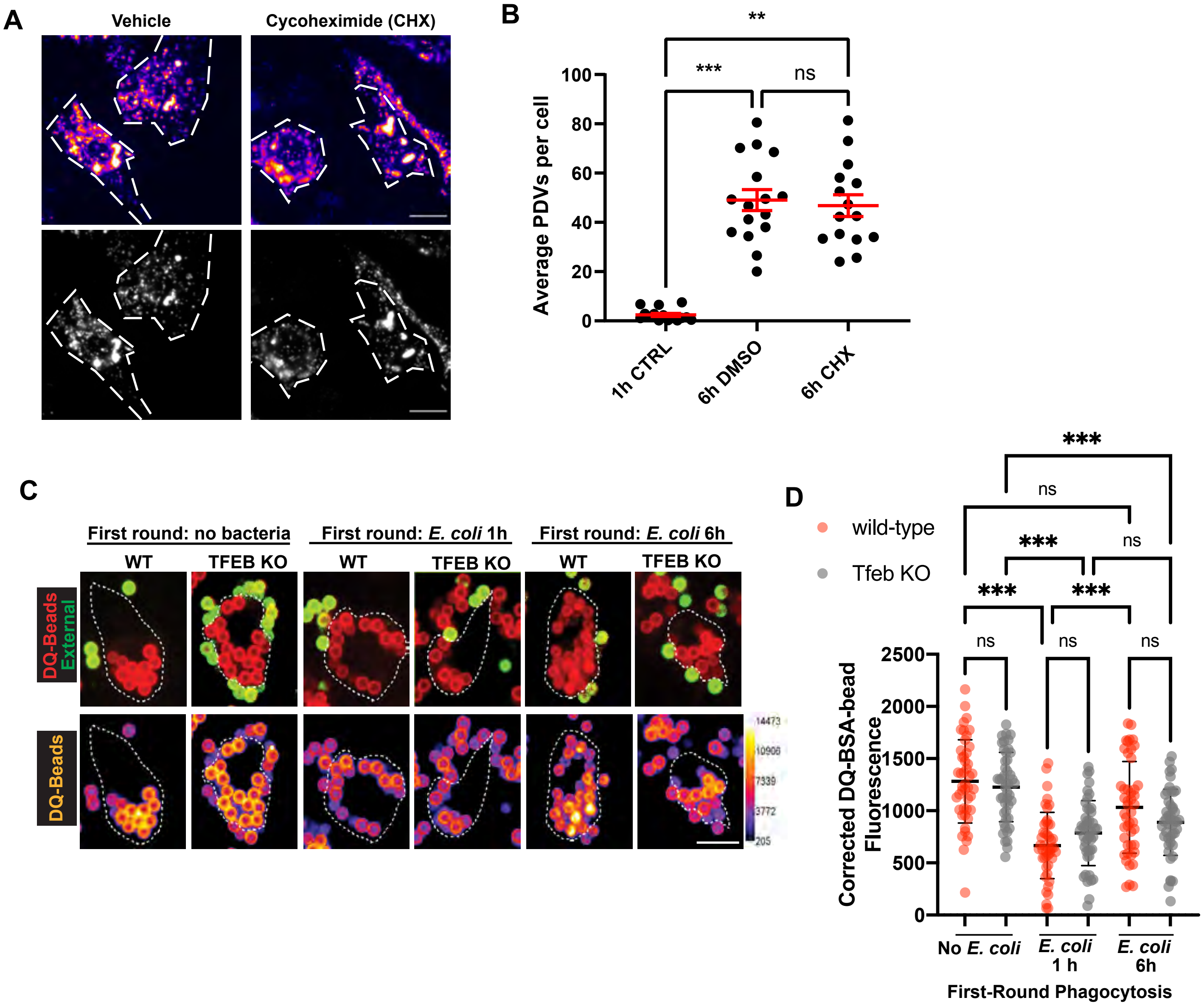


Supplemental Figure S4

### Supplemenatl Figure S5

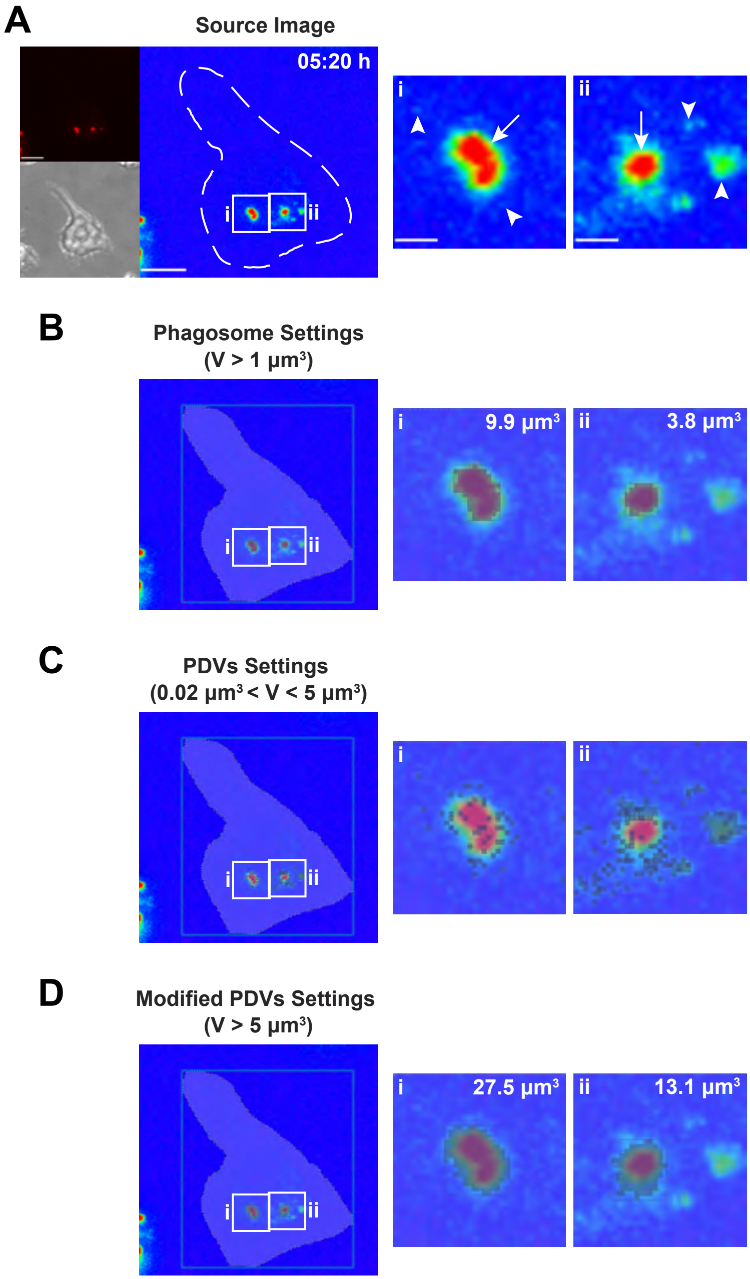
